# Opposing roles of resident and infiltrating immune cells in the defence against *Legionella longbeachae* via IL-18R/IFN-γ/ROS axis

**DOI:** 10.1101/2024.01.31.578217

**Authors:** Lara M. Oberkircher, Victoria M. Scheiding, H. Linda Rafeld, Eric Hanssen, Jan N. Hansen, Markus J. Fleischmann, Nina Kessler, David Pitsch, Dagmar Wachten, Wolfgang Kastenmüller, Andrew S. Brown, Elizabeth L. Hartland, Ian R. van Driel, Garrett Z. Ng, Natalio Garbi

**Author notes:** Corresponding author: Natalio Garbi, Institute of Molecular Medicine and Experimental Immunology, Building 12, Venusberg Campus 1, University Hospital, University of Bonn, D-53127 Bonn, Germany. Authors declare no competing interests.

## Abstract

The immune response against *Legionella longbeachae*, a causative agent of the often-fatal Legionnaires’ pneumonia, is poorly understood. Here we investigated the specific roles of tissue-resident alveolar macrophages (AM) and infiltrating phagocytes during infection with this pathogen. AM were the predominant cell type that phagocytosed bacteria o day after infection. Three and five days after infection, AM numbers were greatly reduced while there was an influx of neutrophils and later monocyte-derived cells (MC) into lung tissue. AM carried greater numbers of viable *L. longbeachae* than neutrophils and MC, which correlated with a higher capacity of *L. longbeachae* to translocate bacterial effector proteins required for bacterial replication into the AM cytosol. Cell ablation experiments demonstrated that AM promoted infection whereas neutrophils and MC were required for efficient bacterial clearance. IL-18 was important for IFN-γ production by IL-18R^+^ NK cells and T cells which, in turn, stimulated ROS-mediated bactericidal activity in neutrophils resulting in restriction of *L. longbeachae* infection. Ciliated epithelial cells also expressed IL-18R but did not play a role in IL-18-mediated *L. longbeachae* clearance. Our results have identified opposing innate functions of tissue-resident and infiltrating immune cells during *L. longbeachae* infection that may be manipulated to improve protective responses.

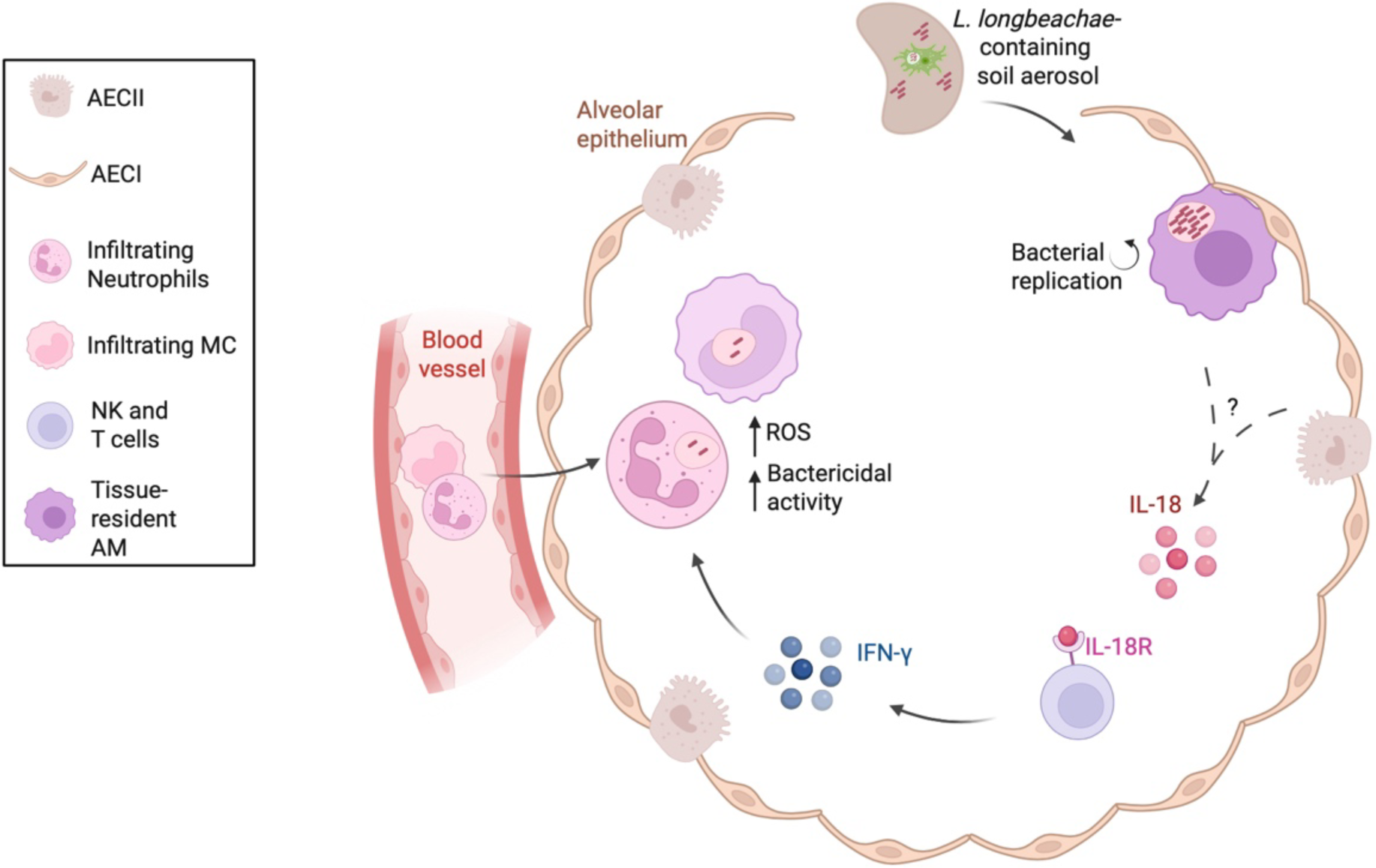

**Highlights:**

1. AM serve as a replicative niche and promote *L. longbeachae* infection.
2. Infiltrating neutrophils and MC kill *L. longbeachae*.
3. IFN-γ from IL-18R^+^ NK and T cells stimulates ROS and bacterial clearance.

## Introduction

The innate immune response is crucial for rapid anti-bacterial defence and prevention of systemic spread, especially from barrier organs such as the lung. For *Legionella* spp., the failure of the innate immune response to control bacterial replication can lead to the development of severe Legionnaires’ disease, an often fatal form of bacterial pneumonia [1]. While the innate immune response to *L. pneumophila* has been extensively studied, much less is known about the innate immune response to *L. longbeachae*, the second most prevalent cause of Legionnaires’ Disease. The increase in *L. longbeachae* infections globally over the past few decades [2] emphasizes the necessity to characterize the cellular and molecular basis of its immune control.

Although *in vitro* studies using murine bone marrow-derived macrophages have suggested that macrophages promote infection by becoming a replicative niche for *L. longbeachae* [3] and *L. pneumophila* [4,5], the relevance of those findings for tissue-resident alveolar macrophages (AM) *in vivo* remains controversial. A recent study demonstrated that depletion of AM increased *L. pneumophila* burden in the lung and worsened disease progression [6]. However, another study found that inducing macrophage apoptosis resulted in an decreased burden of *L. pneumophila* and *L. longbeachae* in the lungs of mice [7]. In addition, monocytes that infiltrate the lung after infection and their progeny such as AM-like cells, inflammatory macrophages, and dendritic cells [8–11], have been shown to be important mediators of defence against *L. pneumophila* and *L. longbeachae* [12]. Whereas it is known that monocyte-derived cells (MC) protect against *L. pneumophila* via production of pro-inflammatory interleukin (IL)-12 [10,12], the mechanisms of protection against *L. longbeachae* are not clear.

The strong influx of neutrophils observed during mouse infections with *L. longbeachae* suggests a protective role of these infiltrating phagocytes [3], similar to that demonstrated for *L. pneumophila* in neutrophil-depleted mice [13]. However, the role of AM and neutrophils as well as other infiltrating leukocytes in the defence against *L. longbeachae* remains unknown.

Flagellin recognition by the NAIP5/NLRC4 inflammasome is critical for defence against *L. pneumophila* infection [12], but it plays no role against the non-flagellated *L. longbeachae* [3]. Therefore, other molecular mechanisms are dominant in the control of *L. longbeachae*. Interferon (IFN)-γ and IL-12 are important for clearance of *L. longbeachae* [3], although the mechanisms of action and the role of other inflammatory cytokines have not been investigated. The successful defence against *L. pneumophila* requires IFN-γ [10,14] as part of a positive feedback loop: MC-derived IL-12 stimulates IFN-γ secretion by NK cells and T cells, which subsequently enhances the bactericidal activity of MC [10]. Understanding the interplay between bactericidal phagocytes and cytokine-producing lymphocytes during infection with *L. longbeachae* might serve as a basis for identifying individuals with a productive innate immune response and who develop severe pneumonia.

Here, we identify a dichotomous role of tissue-resident and infiltrating phagocytes in innate immune defence against *L. longbeachae*. Resident AM readily internalized invading bacteria and supported a high bacterial load, resulting in the establishment of a productive pulmonary infection. In contrast, infiltrating MC and neutrophils restricted *L. longbeachae* viability leading to a reduction in bacterial burden. We further demonstrate that IL-18R expression on lymphocytes rather than on ciliated bronchiolar epithelial cells was required for control of *L. longbeachae* infection via IFN-γ production by NK cells and T cells, which activated ROS production in neutrophils. Our results emphasize a dynamic interplay between infection-promoting tissue-resident AM and protective infiltrating phagocyte populations that may constitute a basis for future development of intervention therapies.

## Material and Methods

### Bacterial strains

The clinical isolate *L. longbeachae* NSW150 (here referred to as *L. lo*) and the recombinant strains *L. longbeachae* carrying pXDC61:BlaM-RalF (*L. lo* BlaM-RalF) and the effector translocation mutant *L. longbeachae ΔdotB* carrying pXDC61: BlaM-RalF (*ΔdotB* BlaM-RalF) were used in this work [15]. All bacteria were grown on buffered charcoal yeast extract (BCYE) agar (Oxoid, Germany) plates at 37 °C for 3 days as previously described [16].

To generate mCherry-expressing *L. longbeachae* (*L. lo*-mCherry), bacteria were grown on BCYE agar at 37 °C for 3 days. Five single colonies were selected, electroporated with 10 ng pON-mCherry plasmid (Addgene, US) in 200 µl ddH_2_O using 2-mm cuvettes at 2.5 kV and 200 Ω for 4 ms. After 3 days on BCYE agar plates, single pink colonies were selected and stored in 50% (v/v) glycerol/ BHI solution at −80 °C until further use.

### Mice

C57BL/6J (The Jackson Laboratory strain #000664), B6.129S4-*Ccr2^tm1/fc^*/J (The Jackson Laboratory strain #004999, referred to here as *Ccr2^−/−^*) [17], B6.129S-*Cybb^tm1Din^*/J (The Jackson Laboratory strain #002365, *gp91phox^−/−^*) [18], B6.129P2-*il18r1^tm1Aki^*/J (The Jackson Laboratory strain #004131, *Il18r1^−/−^*) [19], B6.IL18R^TOM^ (IL-18R reporter mice expressing tdTomato under the endogenous *Il18r1* promoter, referred to as IL-18R tdTomato) [20], B6.Cg-*Commd10^Tg(Vav1-icre)A2Kio^*/J (The Jackson Laboratory strain #008610, *Vav1*^iCre/wt^) [21,22], C57BL/6J-Tg(Nkx2-1-cre)2Sand/J (The Jackson Laboratory strain #008661, *Nkx2.1*^cre/wt^) [23], B6.Cg-*Rag2^tm1.1Cgn^*/J (The Jackson Laboratory strain #008449, *Rag2^−/−^*) [24], and B6.129S7-*Ifng^tm1Ts^*/J (The Jackson Laboratory strain #002287, *Ifng^−/−^*) [25] mice were used. Mice were bred and kept under specific-pathogen-free conditions at the animal facilities of the University Hospital Bonn. In experiments on bacterial effector translocation and IFN-γ deficiency, mice were bred and kept under specific-pathogen-free conditions at the animal facilities of the Bio21 Molecular Science and Biotechnology Institute. Experimental mice were sex- and age-matched (8–12 weeks old). All animal experiments were approved by the Landesamt für Natur-, Umwelt- und Verbraucherschutz, NRW, Germany, or by the University of Melbourne Animal Ethics Committee, Australia.

### Infection of mice with *L. longbeachae* and CFU quantification in lungs

Mice were infected intranasally (i.n.) as previously described [6]. In brief, several bacterial colonies were collected in DPBS after 3 days of growth on BCYE agar at 37°C. Bacterial suspension was initially adjusted in DPBS at an OD_600nm_ of 1.0, amounting to 10^9^ CFU / ml. Mice were administered 50 µl of the indicated inoculum i.n. under isoflurane-induced anaesthesia. The concentration of the infection inoculum was confirmed by retrospective determination of CFU.

*L. longbeachae* CFU in lung homogenates were determined at the indicated time points after 3 days of growth on BCYE agar at 37 °C.

*L. lo*-mCherry CFU were quantified in the indicated phagocytes after lysing FACS-sorted cells in 0.1% Tween-20 for 30min on ice and growing bacteria as described for lung samples. We sorted the total population of the indicated phagocyte subpopulation containing mCherry^+^ and mCherry^−^ cells to obtain enough AM cells for accurate analysis.

### *In vivo* specific cell depletion

Neutrophils were depleted as previously described [26]. In brief, 200 µg anti-Ly6G mAb (clone 1A8, rat IgG2a, BioXcell, US) were applied daily intraperitoneally (i.p.), and every second day in combination with anti-rat IgG2a mAb (clone MAR18.5, BioXcell, US), starting one day prior to infection. Rat IgG2a (clone 2A3, BioXcell, US) was used as isotype control for 1A8.

NK1.1^+^ cells, including NK and NKT cells, were depleted as previously described [27]. Briefly, 200 µg anti-NK1.1 mAb (clone PK136, BioXcell, US) was administered i.p. on day −1, 0, and 1. Infection was performed on day 0.

AM were depleted as previously described [6]. In brief, 50 µl of clodronate-loaded liposomes or PBS-control liposomes (Liposoma B.V., Netherlands) were administered intratracheally (i.t.) one day prior to infection.

### Flow cytometry and cell sorting

Extravascular infiltrating leukocytes in the lung were identified as previously described [28]. Briefly, mice were administered 5 µg Alexa Fluor (AF) 488-labelled anti-CD45.2 antibody (BioLegend, US) intravenously (i.v.) 5 min prior to killing. Lungs were perfused with PBS, sampled and digested in 1 mg/ ml Collagenase IV (Sigma-Aldrich, US) and 100 U/ ml DNase I (Sigma-Aldrich, US) for 30 min at 37 °C. Single-cell suspensions were stained with antibodies listed in **Table S1** and acquired with a LSR Fortessa flow cytometer (BD Bioscience, US). For cell-associated autofluorescence to be removed in the indicated experiments, cell suspensions were acquired with a 7-laser ID7000 spectral flow cytometer (SONY, Japan). Dead cells and doublets were excluded with Fixable Viability Dye eF780 (eBioscience, US) and standard procedures, respectively. Viable cells were analysed using FlowJo V10.8.1. Uniform Manifold Approximation and Projection (UMAP)-based dimensionality reduction was performed using the R-based FlowJo plugin UMAP V3.1.

The indicated phagocyte subpopulations were sorted from single-cell lung suspensions on a FACSAria Fusion (BD Bioscience US) reaching a post-sort purity of > 97% viable phagocytes.

### Imaging flow cytometry

For imaging flow cytometry single-cell suspensions were prepared as described in the section *Flow cytometry and cell sorting*. Hoechst33528 (Sigma-Aldrich, US) was used to exclude dead cells. Hoechst^−^CD45^+^ cells were sorted and then analysed using an ImageStream X MarkII (Amnis) imaging flow cytometer.

### Cytokine profiling from lung homogenate

Cytokines were quantified from 1 ml right lung homogenate using the ProcartaPlex Mouse 12-Plex Mix&Match kit (Thermo Fisher Scientific) according to the manufacturer’s instructions. Data were analysed using the online ProcartaPlex analysis software (Thermo Fisher Scientific).

### Assay for cytosolic translocation of *L. longbeachae* effectors

Detection of BlaM-RalF translocation into mouse cells infected with *L. longbeachae* or the *ΔdotB* mutant carrying pXDC61:BlaM-RalF was performed as previously described [15]. One day after lung infection with 2.5 x 10^5^ CFU *L. lo* BlaM-RalF, or *ΔdotB* BlaM-RalF the BlaM substrate CCF2-AM was added to lung single-cell suspensions and immune cells were analysed by flow cytometry. Translocation of BlaM-RalF into the host cell cytosol results in a fluorescence shift of the CCF2-AM emission maximum from 530 nm (native CCF2-AM) to 460 nm (BlaM-cleaved CCF2-AM), thereby identifying cells supporting translocation of *L. longbeachae* effector molecules.

### Confocal microscopy

Vibratome sections of lungs 2 days after infection were analysed as previously described [9]. Briefly, mice were humanely killed by xylazine/ ketamine overdose and lungs inflated with 1 ml of 2% low melting temperature agarose (Promega, US) at 37 °C. After agarose polymerization, 100-µm lung sections were cut using a VT1000S vibrating-blade microtome (10 Hz and 5–7 µm/ s), fixed using Antigenfix (Diapath, US) for 30 min at 4 °C, and stained overnight at 4 °C with fluorescently labelled antibodies listed in **Table S2**. Sections were mounted in 0.1µg/ml DAPI (BioLegend, US). Imaging was performed with a Zeiss LSM 710 confocal microscope using the spectral lambda mode and linear unmixing to remove autofluorescence, or with a Leica Sp8 Airyscan confocal microscope where indicated. Images were analysed using Imaris software (Bitplane, Switzerland) or ImageJ/ Fiji (v2.8.0/1.53t).

### Correlative Light and Electron Microscopy

After imaging with confocal the samples were post fixed 1h in OsO4 1% in 0.1M Sodium cacodylate, rinse 5×1min in H2O, dehydrated in a series of ethanol then acetone and substituted in resin (Procure 812). The resin was then polymerised between two Aclar films in order to flatten the section. The resulting thin epoxy slab was then glued to a naked epoxy block, trimmed and sent for microCT to monitor the flatness and the high in the resin block. Following analysis of the microCT data the tissue slice was sectioned to 70 nm, stained with Uranyl acetate and Reynold’s lead and observed on a Teneo scanning electron microscope (FEI, Eindovhen, NL) with the backscattered detector. Preliminary alignment was done at low magnification after flipping the fluorescence data image. Alignment of the fluorescence data and the electron microscopy was done using midas (IMOD package, Mastronarde, D.N. (1997) Dual-axis tomography: an approach with alignment methods that preserve resolution. J. Struct. Biol. 120:343-35) using nuclei and nucleoli as fiducial for alignments. The Z average of the fluorescent data was used to align both datasets as the ultrathin EM section was not exactly parallel to the surface of the tissue slice (∼1-3 degres variation in x and y).

### Statistical analysis

Statistical analyses were performed using Prism software v9.4.1 (GraphPad, US). Student’s *t* test for parametric results or the non-parametric Mann–Whitney test was performed to compare two groups. Comparison between three or more groups was performed by one-way ANOVA followed by multiple comparisons according to Tukey’s test for parametric data or the Kruskal–Wallis test followed by Dunn’s multiple comparisons test for non-parametric data. The Mantel–Cox test was performed for mouse survival data. Unless otherwise stated, data are expressed as mean + SD. Differences were considered statistically significant at P < 0.05.

## Results

### Concomitant decrease of alveolar macrophages and influx of circulating immune cells into the lung during *L. longbeachae* infection

Following intranasal (i.n.) inoculation with a sublethal dose, *L. longbeachae* induced a productive infection in mice as demonstrated by a significant weight loss (**Fig 1a**) and increased bacterial load in the lung above the initial infection dose of 10^4^ CFU (**Fig 1b**). Bacterial CFU peaked on days 3–5 after infection and rapidly decreased over the subsequent 5 days. At the peak of infection, we detected various inflammatory cytokines in the lung, including neutrophil-recruiting and neutrophil-stimulating cytokines, such as G-CSF, TNFα, IL-1β and IFN-γ (**Fig 1c**).

**Figure 1:**
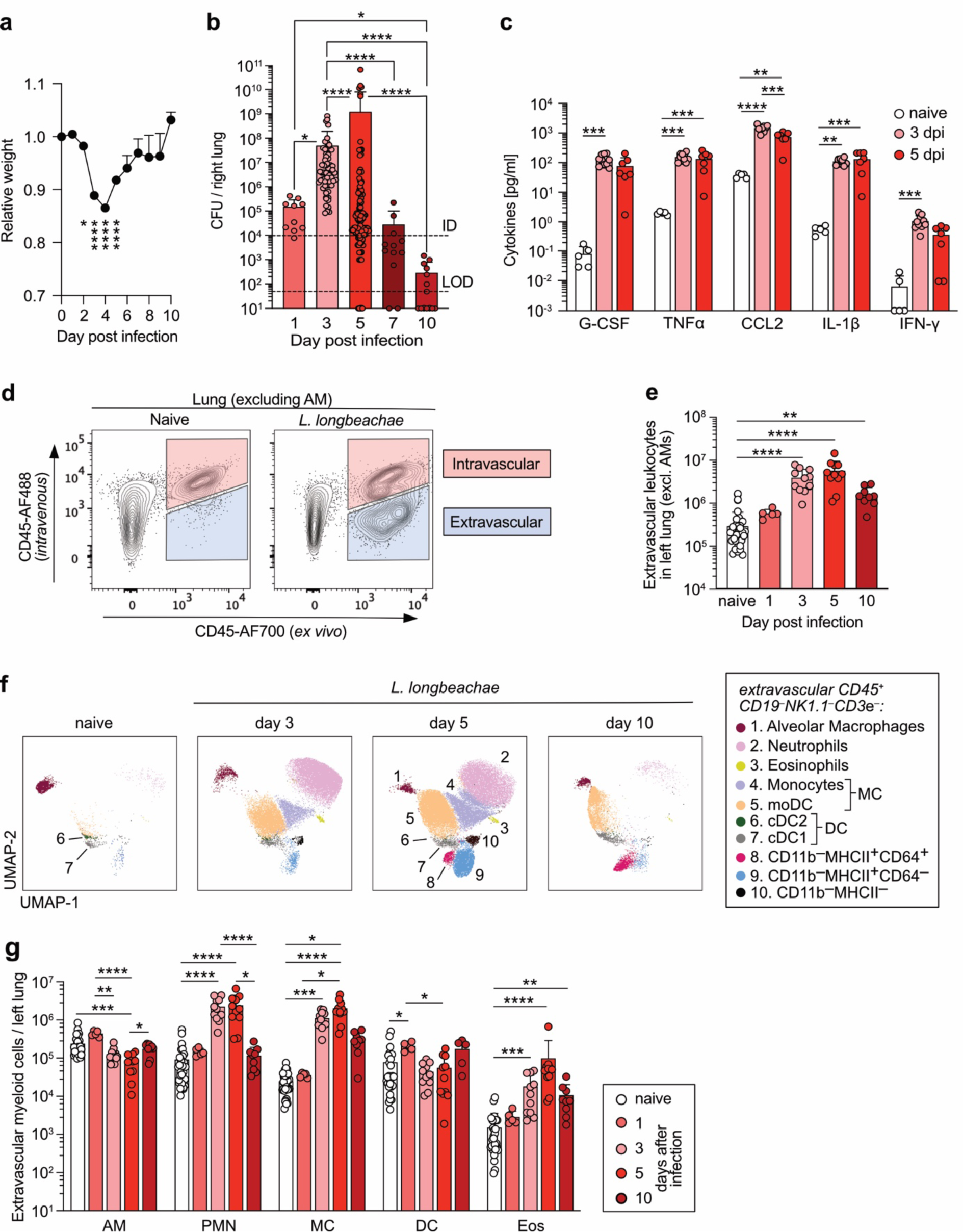
Circulating immune cells infiltrate the lung during infection with *L. longbeachae* while alveolar macrophage numbers are reduced. C57BL/6J mice were intranasally infected with 10^4^ *L. lo* CFU. **(a)** Kinetics of relative mouse body weight during infection. **(b)** Bacterial load in the lung at the indicated time. LOD = limit of detection; ID = infection dose. **(c)** Cytokine quantification in right lung homogenates at the indicated time after infection. Uninfected mice were used as naïve controls. Dpi, days post infection. **(d)** Representative flow cytometry plots of lung cells in naïve mice (left panel) and 3 days after infection (right panel). Red gate indicates intravascular leukocytes; blue gate indicates extravascular leukocytes that have infiltrated the lung. **(e)** Quantification of extravascular leukocytes as identified in (d) at the indicated time points. AM were excluded to enrich for infiltrating leukocytes. **(f)** Flow cytometry UMAP plots of extravascular CD19^−^ NK1.1^−^ CD3ε^−^ myeloid cells and AM at the indicated time in the lung of 5 concatenated mice per group. Gating strategy to identify specific immune cell populations is shown in Supplementary Fig 1. moDC, monocyte-derived dendritic cells, cDC, conventional dendritic cells. **(g)** Quantification of indicated immune cells from (f). Data are shown as mean +SD. Data are pooled from 2 to 3 experiments, or 10 experiments (day 3 and 5 after infection in (a, b)). Symbols represent single mice. Statistical differences were analysed by the Kruskal–Wallis test with Dunn’s post-test and depicted as * P < 0.05, **P < 0.01, *** P < 0.001, **** P < 0.0001.

We investigated the immune cell composition in the lung during bacterial infection. By administering AF488-labelled anti-CD45 i.v., we were able to accurately quantify immune cells that had extravasated into the lung tissue (**Fig 1d**). Infiltration of leukocytes peaked 3–5 days following infection and decreased again on day 10 (**Fig 1e**), which correlated with CFU counts. We performed UMAP-based dimensionality reduction of flow cytometry data to visualize dynamics of different myeloid cell types during infection (**Fig 1f, Supplementary Fig 1**). The immune cell infiltrate from day 3 onward infection was predominantly composed of neutrophils and MC (monocytes and monocyte-derived dendritic cells [11]), in kinetics that positively correlated with CFU (**Fig 1g**). Lung-resident AM, on the other hand, decreased in numbers when bacterial CFU was highest and returned to normal levels by day 10 after infection, thus negatively correlating with bacterial burden (**Fig 1g**).

### A stable *L. longbeachae*-mCherry reporter strain identifies bacteria associated mostly with immune cells

We generated a strain of *L. longbeachae* that expresses the fluorescent protein mCherry (hereafter referred to as *L. lo*-mCherry) to identify cells that associated with *L. longbeachae*. Bacterial colonies showed uniform mCherry expression after 9 days in serial bacterial culture as analysed in 10 randomly picked clones, indicating stable reporter expression *in vitro* (**Fig 2a**). Next, we investigated the stability of mCherry expression during *in vivo* infection. Over 97% of *L. longbeachae* colonies isolated from lungs of infected mice were mCherry^+^ 5 days after infection (**Fig 2b**), demonstrating stable mCherry expression *in vivo*.

**Figure 2:**
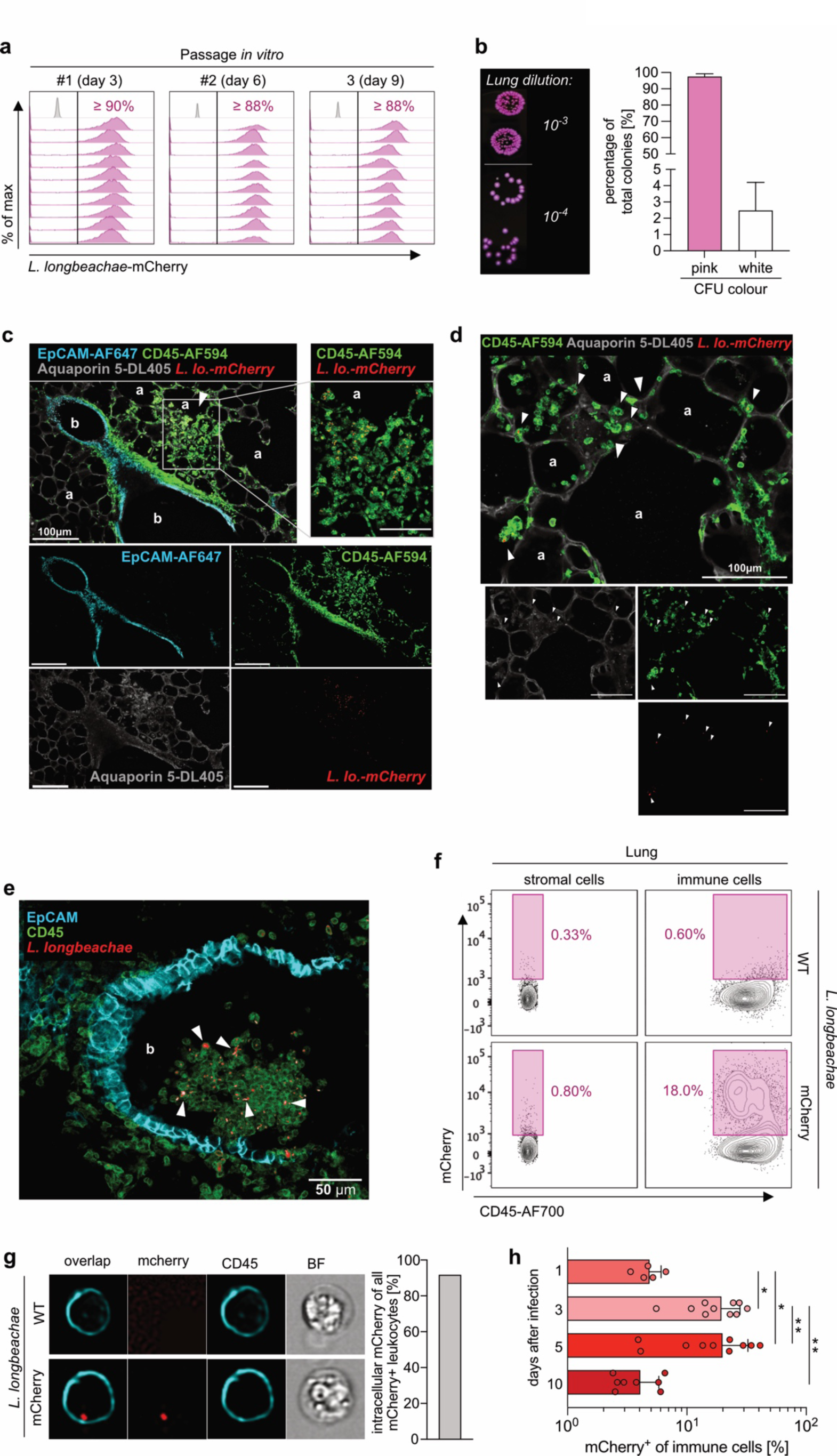
A stable fluorescent *L. longbeachae*-mCherry reporter revealed bacteria mainly located in immune cells during pulmonary infection. Transformed *L. longbeachae* constitutively expressing mCherry (*L. lo*-mCherry) were grown on BCYE agar plates at 37 °C for 3 days before analysis. **(a)** Flow cytometry histograms showing distribution of mCherry expression by bacteria in 10 randomly picked colonies at the indicated passage and time point. Numbers indicate percentage of mCherry^+^ bacteria. **(b)** Representative picture of bacterial colonies grown on BCYE agar from lung homogenate 5 days after infection (left panel), and quantification of mCherry^+^ colonies (right panel). Left panel shows duplicate colonies at the indicated dilution of lung homogenate. A total of 4,400 bacterial colonies derived from 44 mice were quantified in four independent infections. **(c)-(e)** Representative confocal images from lung vibratome slices 5 days after infection of C57BL/6J mice with 10^4^ *L. lo*-mCherry CFU intranasally. a = Aquaporin5^+^ alveoli; b = EpCAM^+^ bronchiole; arrowheads indicate close localization of *L. longbeachae* and CD45^+^ leukocytes. **(f)** Representative flow cytometry plots of stromal cells (left panels) and leukocytes (right panels) showing percentage of cells with *L. lo*-mCherry 3 days after infection of C57BL/6J mice with 10^4^ CFU of the indicated bacteria. **(g)** Representative ImageStream pictures showing uptake of *L. lo*-mCherry by CD45^+^ leukocytes and quantification in 500 cells (right panel) 3 days after infection of C57BL/6J mice with 10^4^ *L. lo*-mCherry CFU. **(h)** Percentage of *L. lo*-mCherry^+^ cells within CD45^−^ stroma and CD45^+^ immune cells 3 days after infection of C57BL/6J mice with 10^4^ CFU. Mice infected with WT *L. longbeachae* served as mCherry^−^ controls. Data are pooled from two experiments with each symbol representing individual mice (h). Data are shown as mean +SD and statistical differences in (h) analysed by repeated measures one-way ANOVA with Tukey’s post-test. * P < 0.05, **P < 0.01.

To identify areas where infection of *L. longbeachae* was established within the lung, we used spectral confocal microscopy of vibratome slices stained with anti-CD45, anti-EpCAM, and anti-aquaporin 5 antibodies. *L. longbeachae* was commonly found together with CD45^+^ leukocytes either as large clusters (**Fig 2c**) or dispersed (**Fig 2d**) throughout aquaporin 5^+^ alveolar areas. In addition, some clusters containing leukocytes and bacteria were present in the bronchiolar lumen suggesting discharge of infected material (**Fig 2e**).

Using conventional and imaging flow cytometry, we demonstrated that bacteria were mainly associated with leukocytes (**Fig 2f, g**, respectively). The frequency of mCherry^+^ leukocytes peaked 3–5 days after infection (**Fig 2h**), positively correlating with bacterial burden in the lung (**Fig 1b**). These data demonstrate the utility of a *L. longbeachae*-mCherry reporter strain with high fluorescence intensity to study bacterial immune cell interactions and here was used to identify leukocytes as the main cell population associated with *L. longbeachae* by microscopy and flow cytometry.

### Neutrophils show superior bactericidal activity against *L. longbeachae* compared to alveolar macrophages

We used *L. lo*-mCherry to investigate the role of different resident and infiltrating phagocyte populations during infection (**Supplementary Fig 2**). We identified mCherry^+^ AM, neutrophils, and MC 3 days after infection (**Fig 3a).** UMAP dimensionality reduction of flow cytometry data (**Fig 3b**) and quantification (**Fig 3c**) of phagocytes in the mCherry^+^ population revealed that early during infection (day 1) *L. longbeachae* were mostly associated with AM. The bacteria associated with neutrophils and MC at later time points, with neutrophils being predominant on day 3 and MC on day 10 (**Fig 3c**). We did not detect significant mCherry positivity in other myeloid cell populations (**Fig 3b**).

**Figure 3:**
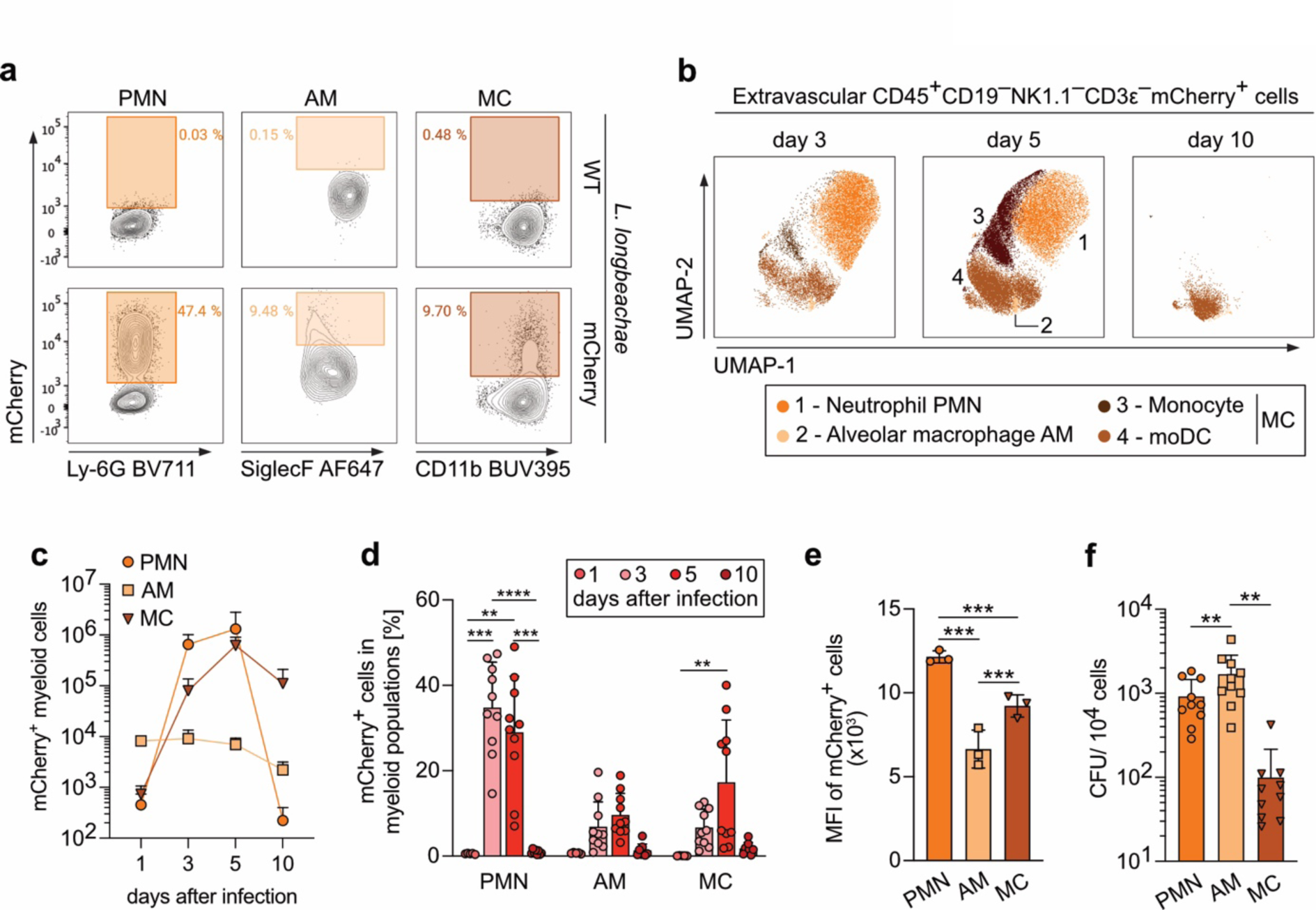
Neutrophils and MC show the highest association with L. longbeachae whereas AM support bacterial viability. C57BL/6J mice were intranasally infected with 10^4^ *L. lo*-mCherry CFU. **(a)** Flow cytometry plots of the indicated myeloid cells 3 days after infection. Gating strategy to identify specific immune cell populations is shown in **Supplementary Fig S2**. **(b)** Flow cytometry UMAP plots of extravascular mCherry^+^ CD19^−^ NK1.1^−^ CD3ε^−^ myeloid cells at different time points after infection. Gating strategy as in (a). **(c)** Quantification of indicated immune cells from (b). **(d)** Percentage of mCherry^+^ cells within the indicated phagocyte population at the indicated times after infection. Cells from mice infected with WT *L. lo* served as mCherry^−^ controls. **(e)** mCherry MFI (mean fluorescence intensity) in the indicated mCherry^+^ cell population 3 days after infection. **(f)** Number of viable bacteria (CFU) per 10^4^ cells of indicated cell population sorted 3 days after infection. Data are pooled from two experiments (b, c, d, f) or performed once (e). Symbols represent single mice. In (b), data are concatenated from 5 mice per time point. In (c) data ae pooled from two experiments with a total of 10 mice. Data are shown as mean +SD. Data were tested for normality and then for statistical differences by the Kruskal–Wallis test with Dunn’s multiple comparison test or one-way ANOVA with Tukey’s multiple comparison test as appropriate. * P < 0.05, ** P < 0.01, *** P < 0.001, **** P < 0.0001.

We then investigated which phagocytes had the strongest association with *L. longbeachae* by quantifying the proportion of mCherry^+^ cells within each phagocyte population. During the peak of infection (day 3 and 5), a higher percentage of neutrophils were positive for *L. lo*-mCherry than AM and MC (**Fig 3d**).

Having identified phagocytes positive for *L. lo-*mCherry, we next assessed the number of viable bacteria associated with different phagocyte populations. For this, we first quantified *L. longbeachae*-derived mCherry mean fluorescence intensity (MFI) using spectral flow cytometry to eliminate cell-associated autofluorescence, particularly that of AM (**Supplementary Fig S3**). We found the highest mCherry MFI in neutrophils, followed by MC and then AM (**Fig 3e**), suggesting that neutrophils carried more bacteria on a per-cell basis. As both live and dead *L. lo-*mCherry can contribute to mCherry MFI, we next identified where viable bacteria were residing. Individual cell populations were FACS-sorted and the number of viable bacteria per 10^4^ cells quantified by retrospective determination of CFU. Despite AM having the lowest mCherry MFI (**Fig 3d, e**), these phagocytes contained the highest number of viable bacteria 3 days after infection (**Fig 3f**), suggesting that AM acted as the main replicative niche *in vivo*. In contrast, neutrophils and MC showed lower number of viable bacteria compared to AMs (**Fig 3f**) despite showing higher mCherry MFI (**Fig 3d, e**), suggesting PMN and MC were able to efficiently eliminate *L. longbeachae*.

### Alveolar macrophages promote establishment of *L. longbeachae* infection in the lung

The finding indicating that AM likely act as a replicative niche to promote *L. longbeachae* infection, as is known for *L. pneumophila* [4,5] led us to investigate whether *L. longbeachae* preferentially translocates bacterial Dot/Icm type IV secretion system (T4SS) effector proteins into the cytosol of AM, a process that is required to form an intracellular replicative niche in macrophages *in vitro* [5,29]. Using the *L. lo* BlaM-RalF effector translocation reporter strain expressing a fusion protein of the prototypic T4SS effector RalF together with the β-lactamase (BlaM) reporter enzyme [15], we were able to quantify RalF effector translocation into the host cytosol 1 day after infection by staining cells with BlaM-sensitive CCF2 dye. Quantification of the shift in fluorescence emission from 530 nm (intact CCF2) to 460 nm (BlaM-cleaved CCF2) (**Fig 4a**, left panel) reflects the level of RalF translocation. Infection with *ΔdotB* BlaM-RalF, which lacks a functional T4SS was used as a translocation-deficient control (**Fig 4a**, right panel). Following infection with *L. lo* BlaM-RalF, all phagocytes showed cytosolic translocation of BlaM-RalF into the cytosol, albeit to a greater extent in AM compared to neutrophils (**Fig. 4b**), which was consistent with AM fulfilling a function as replicative niche for *L. longbeachae*.

**Figure 4:**
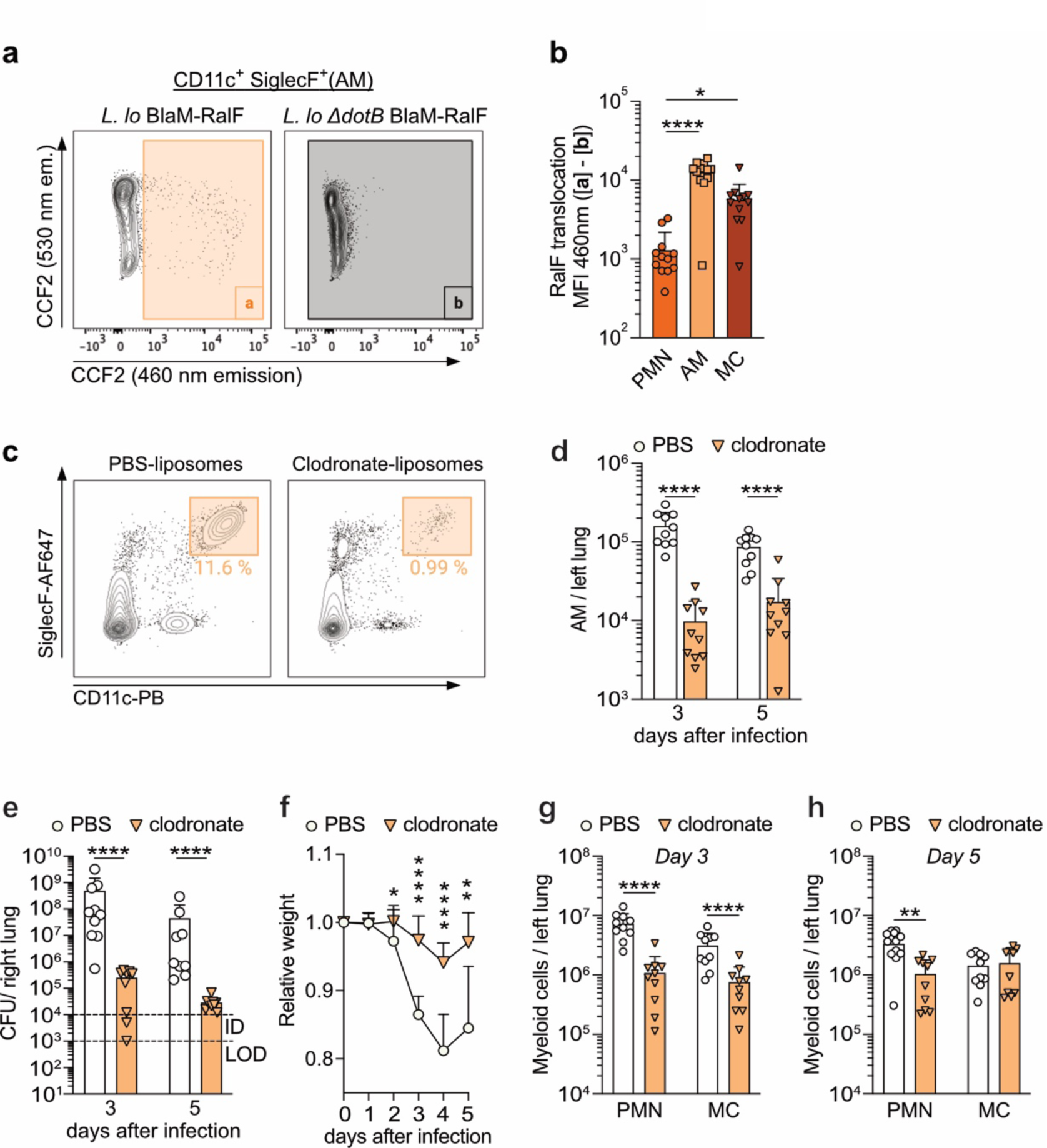
Depletion of alveolar macrophages reduced bacterial burden and weight loss during infection with *L. longbeachae*. **(a), (b)** C57BL/6J mice were intranasally infected with 2.5 x10^5^ CFU *L. lo*. BlaM-RalF or *ΔdotB* BlaM-RalF. **(a)** Representative flow cytometry plots showing the gate to calculate CCF2_460_ MFI resulting from BlaM-RalF translocation (left panel), and background CCF2_460_ MFI (right panel) in AM from mice infected 1 day earlier with the indicated bacterial strain. **(b)** Quantification of RalF translocation in different phagocyte populations *in vivo* as outlined in (a). **(c–h)** C57BL/6J mice were intratracheally administered clodronate- or PBS-loaded liposomes and infected with 10^4^ *L. lo*-mCherry CFU one day later. **(c)** Representative flow cytometry plots showing frequency of AM in the lung 3 days after infection and treated as indicated. **(d)** Quantification of AM at the indicated time and treatment. Kinetics of bacterial load **(e)** and relative mouse weight **(f)** of AM-deficient and AM-competent mice. LOD = limit of detection; ID = infection dose. Quantification of pulmonary myeloid cells at day 3 **(g)** and day 5 **(h)** after infection. Data are pooled from two experiments. Symbols represent single mice. Data are shown as mean +SD. Data were tested for normality and then for statistical differences by the Kruskal–Wallis test with Dunn’s multiple comparison test, Student’s *t* test, or Mann–Whitney test as appropriate. * P< 0.05, ** P < 0.01, *** P < 0.001, **** P < 0.0001.

To directly address the role of AM in promoting bacterial replication during infection, we depleted AM using a single local application of clodronate-loaded liposomes into the lung that resulted in prolonged AM depletion for 21 days during steady state (**Supplementary Fig 4**). When applied one day prior to infection, clodronate liposomes also resulted in approximately 90% reduction in AM numbers both at 3 and 5 days after infection (**Fig 4c, d**). Concomitantly, AM depletion led to a more than 99.9% reduction in bacterial burden in the lungs (**Fig 4e**) and less pronounced weight loss in infected animals (**Fig 4f**). Flow cytometry analysis revealed that AM depletion did not enhance infiltration of neutrophils and MC (**Fig 4g, h**), but rather led to a decrease in their numbers, most likely because of a decreased bacterial load in the absence of AM. Taken together, these results indicate that AM promote *L. longbeachae* infection *in vivo*.

### Neutrophils and monocyte-derived cells are required for restriction of *L. longbeachae* during different stages of infection

To investigate the contribution of neutrophils and MC to *L. longbeachae* defence *in vivo*, we employed knockout mice and cell type-specific depletions to understand the role of each in the innate immune response. *Ccr2^−/−^* mice carrying a defect for monocyte egress from the bone marrow, showed an ∼90% reduction in the number of MC in the lung during *L. longbeachae* infection (**Fig 5a**). Three days after intranasal infection mice there was no difference in bacterial load between WT and *Ccr2^−/−^* mice. However, five days after infection *Ccr2^−/−^* mice showed a 10-fold increase in CFU (**Fig 5b**), indicating that MC play a protective role at a later stage of infection, as previously reported [12].

**Figure 5:**
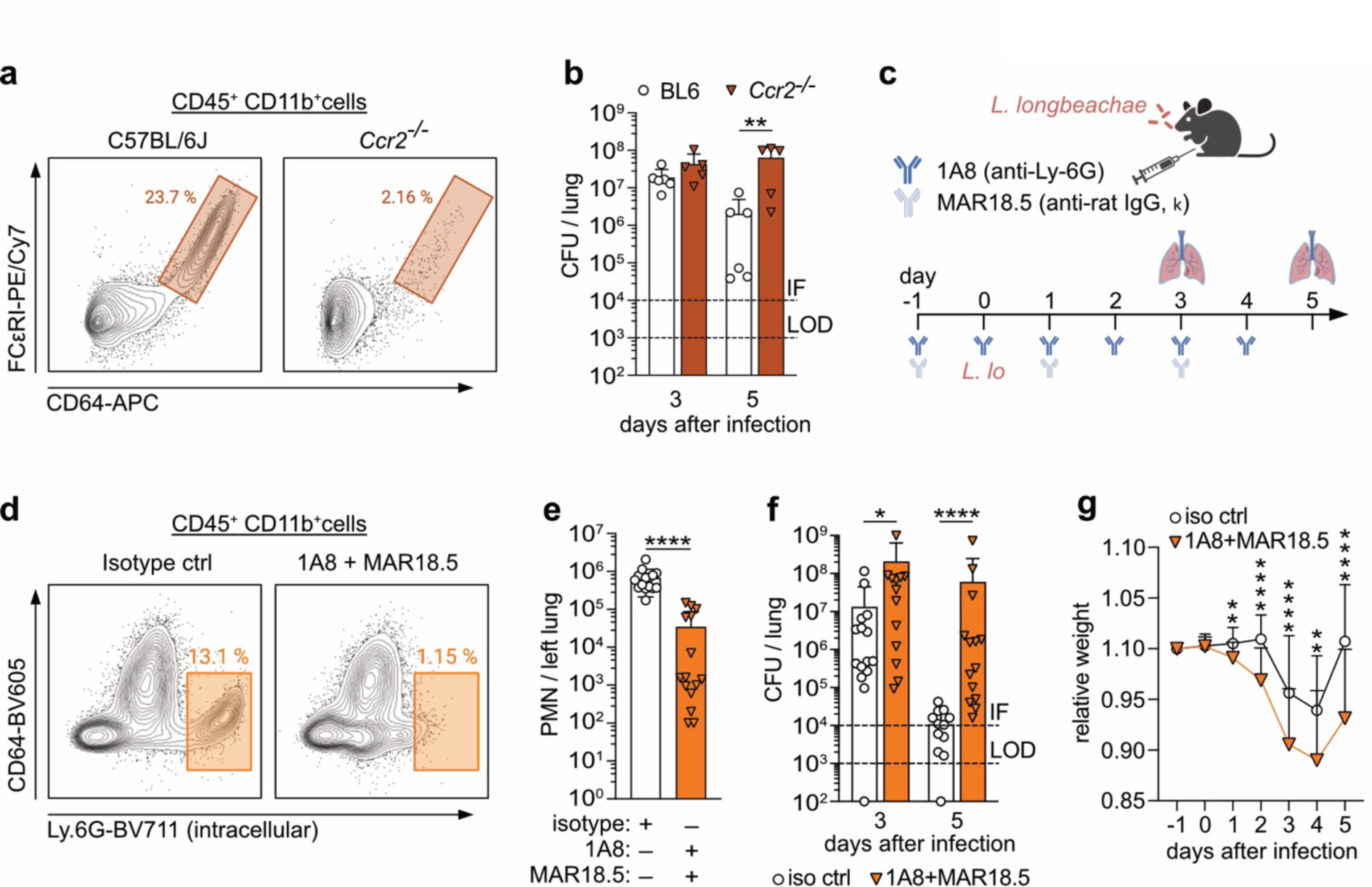
Decreased number of neutrophils and monocyte-derived cells results in impaired bacterial clearance and more pronounced reduction in weight loss. Mice were intranasally infected with 10^4^ *L. longbeachae*. **(a)** Representative flow cytometry plots showing reduced frequency of MC in *Ccr2^−/−^* mice on day 5 after infection. **(b)** Bacterial load in the right lung of the indicated mice. **(c)** Scheme of experimental set-up for neutrophil depletion in (d–g). C57BL/6J mice were intraperitoneally injected with anti-Ly6G (1A8) and secondary anti-rat IgG (MAR18.5) depletion antibody or isotype control starting one day prior to infection. **(d)** Representative flow cytometry plots showing frequency of lung neutrophils intracellularly stained for Ly-6G 5 days after infection. **(e)** Quantification of neutrophils from (d). **(f), (g)** Kinetics of bacterial load in the right lung (f) and relative mouse weight (g) of neutrophil-depleted and neutrophil-nondepleted mice. LOD = limit of detection; ID = infection dose. Data in (b, e, f, g) are pooled from two experiments. Symbols represent single mice. Data are shown as mean +SD. Data were tested for normality and then for statistical differences by Student’s *t* test or Mann–Whitney test as appropriate. * P < 0.05, ** P < 0.01, *** P < 0.001, **** P < 0.0001.

To investigate the function of neutrophils during *L. longbeachae* infection, we followed a combinatorial depletion strategy consisting of antibodies directed against Ly6G (clone 1A8, rIgG2a) and rIgG2a (clone MAR18.5) [26] (**Fig 5c**), to deplete neutrophils prior to infection (**Fig 5d, e**). Neutrophil depletion led to failure to control bacterial burden in the lung as demonstrated by significantly increased CFU counts already on days 3 and 5 after infection (**Fig 5f**), and by more pronounced weight loss (**Fig 5g**). These experiments show both neutrophils and MC were important for restricting bacterial burden in the lung during *L. longbeachae* infection, with neutrophils acting early after infection and MC at later stages.

### IL-18R expression by lymphocytes is required for restriction of *L. longbeachae*

Having established that neutrophils and MC play a protective role during *L. longbeachae* infection, we were interested in the cytokine networks that also mediate resistance to *L. longbeachae*. IL-18 confers protection against certain intracellular bacteria [30–33], and we detected IL-18 in lungs 3 and 5 days after infection with *L. longbeachae* (**Fig 6a**). To study the role of IL-18 in protection against *L. longbeachae*, we infected *Il18r1^−/−^* mice. At 5 days after infection, *Il18r1^−/−^* mice had 1,000-fold higher CFU counts compared to WT mice (**Fig 6b**) and more pronounced reduction in body weight (**Fig 6c**). Consistent with this, *Il18r1^−/−^*neutrophils and MC contained a more than 10-fold higher number of viable *L. longbeachae* than their WT counterparts as indicated by CFU quantification in sorted cells on day 5 after infection (**Fig 6d**).

**Figure 6:**
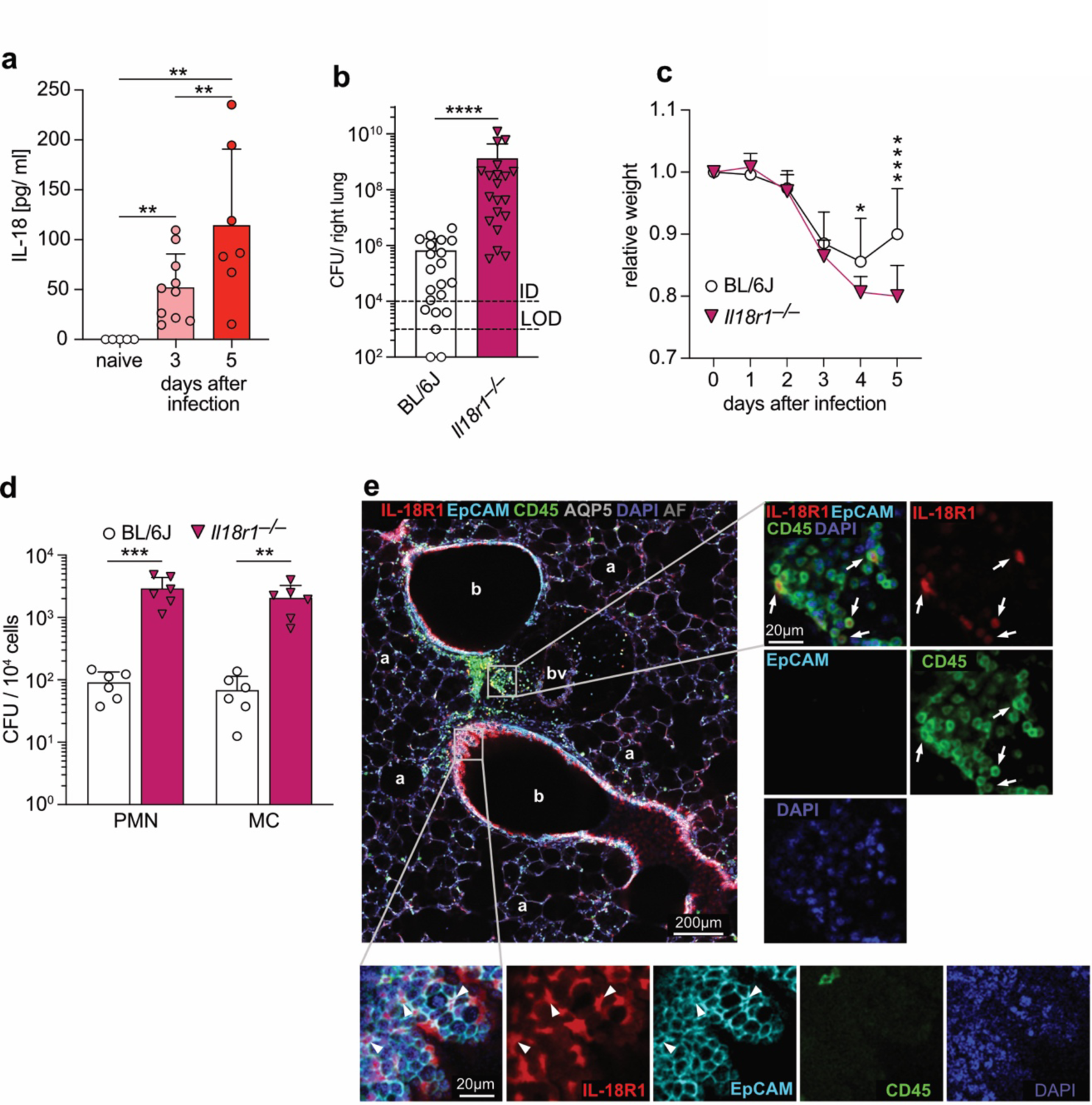
Lack of IL-18R impairs bacterial clearance and increases weight loss. **(a)** Quantification of IL-18 in the right lung homogenate of C57BL/6J at the indicated times after intranasal infection with 10^4^ *Legionella longbeachae* CFU. **(b), (c)** Bacterial load in the right lung 5 days after infection (b) and kinetics of relative body weight of C57BL/6J and *Il18r1^−/−^*mice (c). LOD = limit of detection; ID = infection dose. **(d)** Number of viable bacteria per 10^4^ cells of the indicated total population sorted from the lung 3 days after infection of C57BL/6J and *Il18r1^−/−^* mice. **(e)** Spectral confocal image of vibratome lung slices from IL-181-tdTomato reporter mice 3 days after infection. Insets show magnification of immune cells (right panels) and an EpCAM^+^ bronchiole (bottom panels). Arrows indicate examples of IL-18R1^+^ leukocytes; arrowheads indicate examples of IL-18R1^+^ epithelial cells. a, alveoli; b, bronchiole; bv, blood vessel. Data are pooled from two (a, d) or three (b) experiments; symbols represent single mice. Data in (c) is a representative of five experiments with seven mice. Data are shown as mean +SD. Data were tested for normality and then for statistical differences by Student’s *t* test or Mann–Whitney test as appropriate. * P < 0.05, ** P < 0.01, **** P < 0.0001.

To identify which cells expressed IL-18R in the lung, we used IL-18R-tdTomato mice, in which the *Il18r1* promoter drives tdTomato expression [20]. In immunofluorescence experiments of vibratome sections, IL-18R1^+^ immune cells (CD45^+^ tdTomato^+^) mostly localized within the alveolar space 3 days after infection (**Fig 6e**). In addition, we identified CD45^−^ EpCAM^+^ epithelial cells lining the bronchiolar wall that expressed the IL-18R (**Fig 6e**).

Flow cytometric analysis of cells infiltrating the lung after *L. longbeachae* infection revealed that IL-18R1 was predominantly expressed by NK cells, T cells, and ILC (innate lymphoid cells), with other lymphocytes and myeloid cells having negligible expression (**Fig 7a**), which is consistent with previous reports on IL-18R expression [31]. Within the bronchiolar wall, both confocal microscopy (**Fig 7b, supplementary Video 1**) as well as correlative light and electron microscopy (**Fig 7c**) identified that IL-18R1 was expressed by ciliated epithelial cells (ciEpC) rather than neighbouring uteroglobin^+^ secretory Club cells.

**Figure 7:**
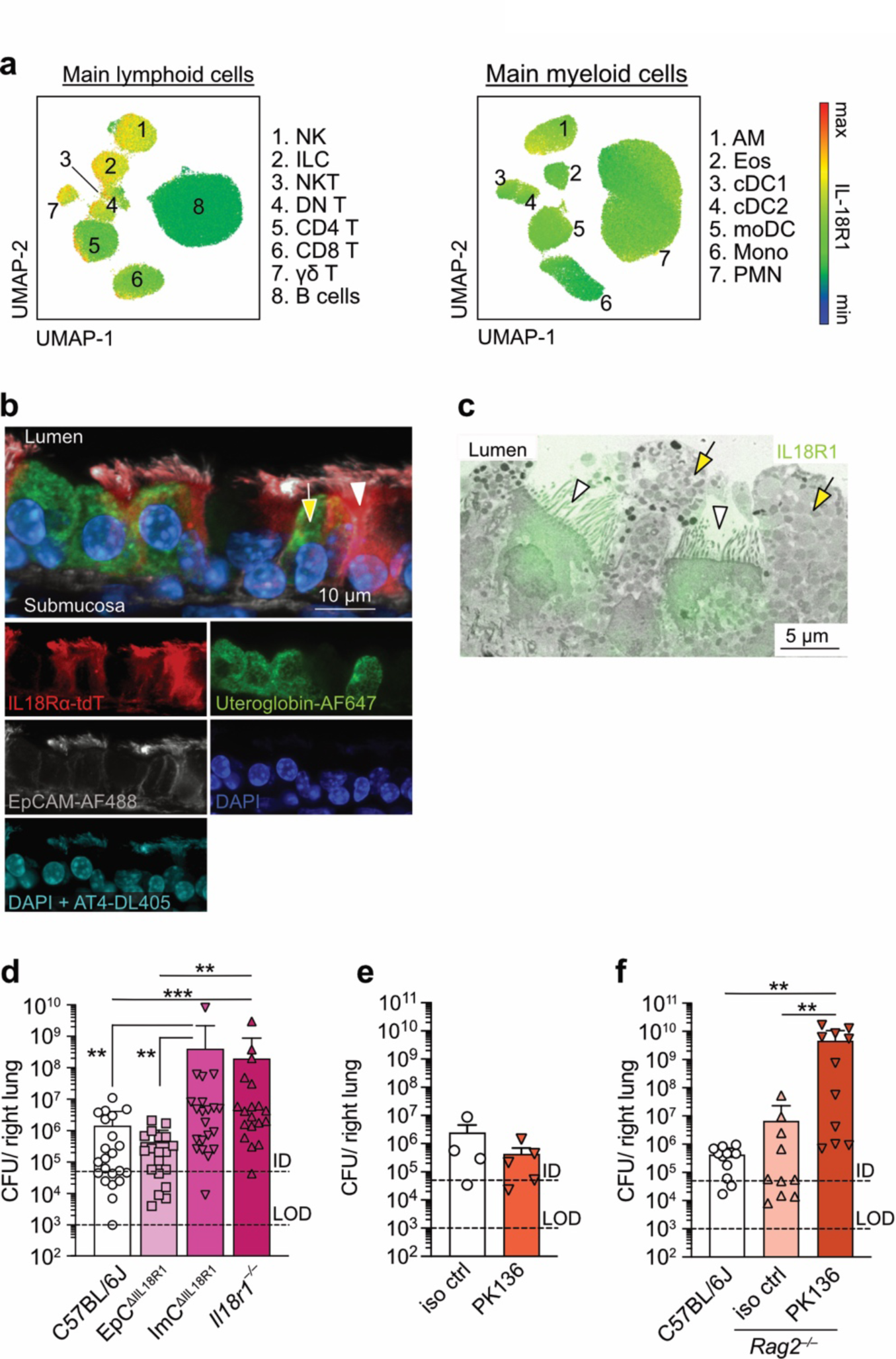
IL-18R expression on either NK cells or T cells rather than on ciliated epithelial cells is essential for defence against *L. longbeachae* in the lung. **(a)** Flow cytometry UMAP plots showing IL-18R1 expression levels by CD45^+^CD11c^−^Ly-6G^−^ lymphoid cells and CD45^+^CD19^−^NK1.1^−^CD3χ^−^ myeloid cells 5 days after infection of IL-18R tdTomato reporter mice with 10^4^ *L. longbeachae* CFU. Gating strategy to identify specific immune cell populations is shown in **Supplementary Fig 1**. **(b)** Representative confocal image of vibratome lung slices of IL-18R1-tdTomato reporter mice showing IL18R1 expression in bronchiolar ciliated (EpCAM^+^ acetylated tubulin IV (AT4)^+^, arrow) and secretory (EpCAM^+^ Uteroglobin^+^, arrowhead) epithelial cells. **Supplementary Video 1** is obtained from the Z stack. **(c)** Correlative light and electron microscopy image of vibratome lung slices of IL-18R1-tdTomato reporter mouse (tdTomato in green) showing ciliated (arrow) and secretory (arrowhead) bronchiolar epithelial cells. **(d)** Bacterial load 5 days after intranasal infection with 5 x 10^4^ *L. lo*-mCherry CFU in the indicated mouse strains. EpC^ΔIL18R^ (*Nkx2.1*^Cre/wt^ x *Il18r1^fl/fl^*); ImC^ΔIL18R^ (*Vav1*^iCre/wt^ x *Il18r1^fl/fl^*. **(e)** Bacterial load as in (d) in C57BL/6J mice treated with depleting anti-NK1.1 antibody (PK136) or mIgG2a isotype control (iso ctrl). **(f)** Bacterial load as in (d) in C57BL/6J or *Rag2^−/−^* mice treated with depleting anti-NK1.1 antibody (PK136) or mIgG2a isotype control (iso ctrl). LOD = limit of detection; ID = infection dose. All data are pooled from two (e) or three (d, f) experiments. Symbols represent single mice. Data are shown as mean +SD. All data were tested for normality and then for statistical differences by the Kruskal–Wallis test with Dunn’s multiple comparison test or Mann–Whitney test as appropriate. * P < 0.05, ** P < 0.01, *** P < 0.001, **** P < 0.0001.

To identify which IL-18R^+^ cell population was required for protective anti-*L. longbeachae* defence, we used conditional knock-out mice where *Il18r1* was deleted either from pulmonary epithelial cells (*Nkx2.1*^Cre/wt^ x *Il18r1^fl/fl^*, termed EpC^ΔIL18R^) or fromimmune cells (*Vav1^iCre/^*^wt^ x *Il18r1^fl/fl^*, termed ImC^ΔIL18R^) (**Supplementary Fig 5**). The bacterial load in *L. longbeachae*-infected ImC^ΔIL18R^ mice was significantly higher than either WT or EpC^ΔIL18R^ mice and comparable to *Il18r1^−/−^* mice, indicating that IL-18R expression in immune cells and not epithelial cells is important for resistance to *L. longbeachae* (**Fig 7d**). Consistent with this finding, IL-18R signalling did not alter ciliary beating frequency or mucus production, two of the main anti-bacterial defence mechanisms employed by EpC (**Supplementary Fig 6**).

To identify which IL-18R^+^ immune cells were involved in *L. longbeachae* defence, we first focused on NK cells and T cells because they infiltrated the lung (**Supplementary Fig 7**) and expressed IL-18R1 during infection (**Fig 7a**). Levels of bacteria in *L. longbeachae*-infected mice that were depleted of NK cells by antibody treatment (**Fig 7e**) as well as in *Rag2^−/−^*mice that lack T and B lymphocytes (**Fig 7f**) were unchanged from control mice. On the other hand, when NK cells were depleted in *Rag2^−/−^* mice, there was a ∼500-fold increased bacterial burden in the lung 5 days after infection (**Fig 7f**, **Supplementary Fig 8**). Given that T cells and not B cells expressed IL-18R (**Fig 7a**), these data suggests a redundant role of NK cells and T cells in protection against *L. longbeachae* infection. ILC also expressed IL-18R1 in the lung (**Fig 7a**) and, although they may further contribute to protection, they could not compensate for the lack of NK and T cells (**Fig 7f**).

### IL-18R-dependent IFN-γ and ROS production promote *L. longbeachae* clearance *in vivo*

Having identified IL-18R^+^ NK cells and T cells as key cell compartments in the defence against *L. longbeachae* infection, we investigated molecular mechanisms behind this protection. IL-18 was initially identified as an IFN-γ-inducing factor [34] and, in line with this, IFN-γ deficiency resulted in an impaired capacity to control *L. longbeachae* infection (**Fig 8a**) and increased bacterial load in neutrophils and MC (**Fig 8b**). Significantly less IFN-γ was found in the lungs of *L. longbeachae*-infected *Il18r1^−/−^* mice than in those of WT mice (**Fig 8c**), consistent with the function of IL-18 to induce IFN-γ [34].

**Figure 8:**
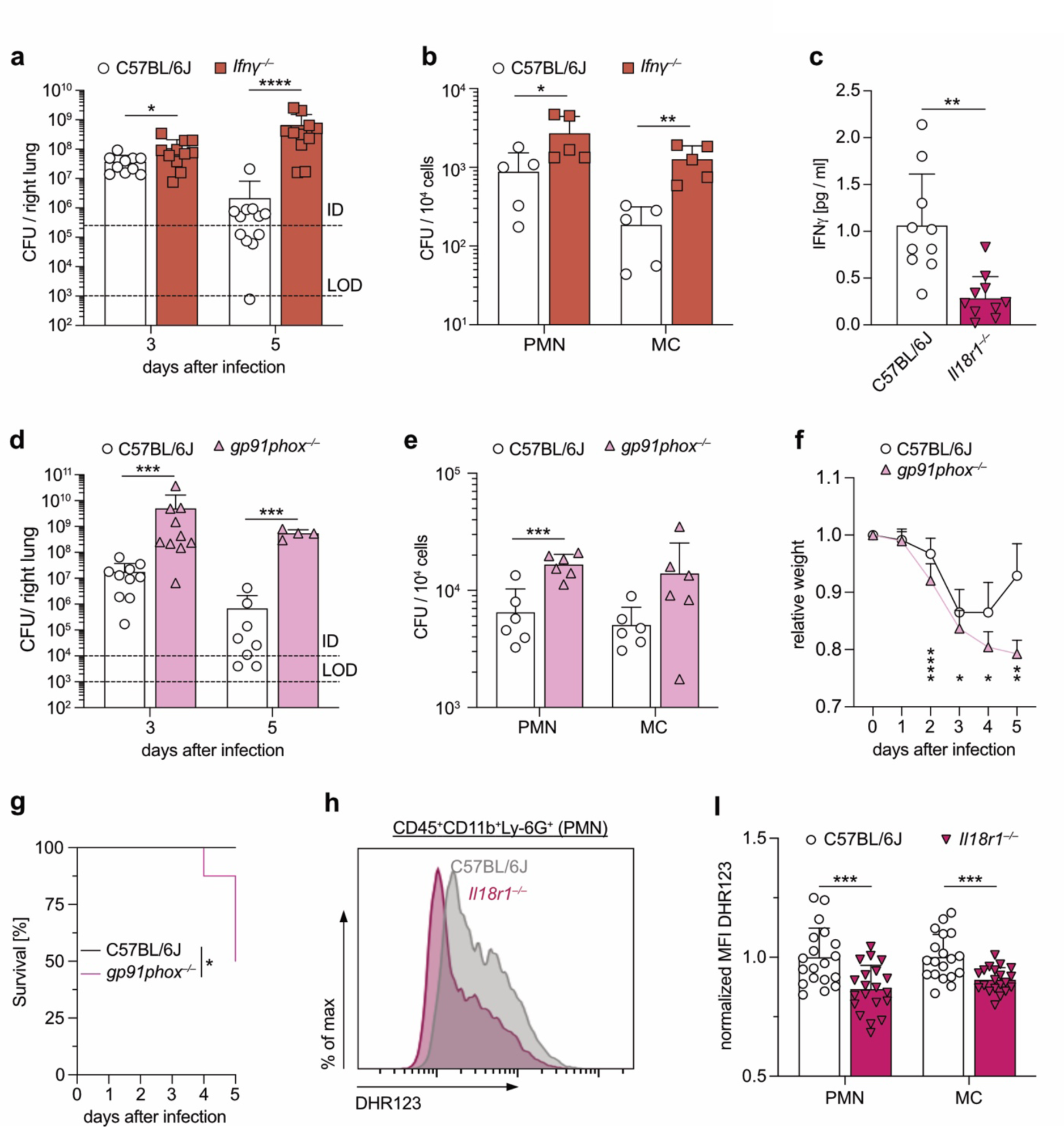
IL-18R1-dependent defence against *L. longbeachae* involves IFN-γ and NADPH oxidase-dependent ROS. Mice were intranasally infected with 2.5×10^5^ (a, b) or 10^4^ (c-i) CFU *L. lo*-mCherry. **(a)** Bacterial load in the right lung at the indicated time after infection of C57BL/6J and *Ifng^−/−^*mice. LOD = limit of detection; ID = infection dose. **(b)** Number of viable bacteria per 10^4^ cells of the indicated total population sorted 3 days after infection of C57BL/6J and *Ifng^−/−^*mice. **(c)** Quantification of IFN-γ in left lung homogenate of indicated mice 5 days after infection. **(d)** Kinetics of bacterial load in the right lung of the indicated mice. **(e-g)** Bacterial CFU per 10^4^ of the indicated cells sorted 3 days after infection from C57BL/6J and *gp91phox^−/−^* mice (e), relative mouse weight (f), and survival (g) of infected C57BL/6J and *gp91phox^−/−^* mice. **(h, i)** Representative flow cytometry histogram (h) and quantification (i) of spontaneous ROS production (DHR123) by neutrophils from the lungs of indicated mice 3 days after infection. Data are pooled from two to three experiments. Symbols represent single mice. As indicated, 50% of *gp91phox^−/−^* mice died by day 5 (g) and thus only remaining mice were analysed on day 5 in (d) and (f). Data were tested for normality and then for statistical differences by Student’s *t* test, Mann–Whitney test, or Mantel–Cox test as appropriate. * P < 0.05; ** P < 0.01; *** < 0.001, **** P < 0.0001.

IFN-γ promotes expression of microbiocidal NADPH oxidase-dependent reactive oxygen species (ROS) by phagocytes [35], and thus we sought to investigate whether NADPH oxidase mediated defence explained the capacity of neutrophils to restrict *L. longbeachae* infection in vivo. Indeed, infection of *gp91phox^−/−^*mice showed that NADPH oxidase was required to control *L. longbeachae* infection in the lung (**Fig 8d**), and in neutrophils (**Fig 8e**), with *gp91phox^−/−^* mice showing greater body weight loss (**Fig 8f**) and decreased survival (**Fig 8g**) compared to WT mice. Finally, we wee able to demonstrate that IL-18R expression promoted production of ROS in neutrophils and MC during *L. longbeachae* infection (**Fig 8h, i**).

Taken together, our results suggest an important role for the classic pathway of IL-18-dependent defence [31,36] during infection with *L. longbeachae* involving IFN-γ secretion from IL-18R^+^ NK cells and T cells that promote downstream ROS production and thereby bactericidal function of neutrophils and MC.

## Discussion

Using a murine model of pulmonary infection with *L. longbeachae*, we have identified a differential contribution of lung-resident and infiltrating phagocytes to disease progression. AM favoured translocation of bacterial T4SS effectors and promoted establishment of infection whereas infiltrating neutrophils and MC were required for host protection. Neutrophils were rapidly recruited into the lung after infection while MC showed more delayed kinetics. Our data demonstrated a clear role of IL-18 in the defence against *L. longbeachae* and suggested that IL-18R-dependent IFN-γ secretion by either NK cells or T cells is sufficient for efficient ROS production by neutrophils.

AM make up a large proportion of tissue-resident immune cells in the lung during homeostasis [37–39] and are among the first immune cells encountering *L. longbeachae* after inhalation. However, we observed that only a low percentage of AM took up invading *L. longbeachae*, most likely because of dispersed infection foci throughout alveoli early during infection. Although it is generally accepted that the overall effect of AM during *L. pneumophila* challenge is to promote infection by acting as a bacterial replicative niche [4,40,41], recent data dispute this role by demonstrating that AM have a net protective function *in vivo* [6]. In contrast to *L. pneumophila*, our data showed that AM depletion led to strongly reduced *L. longbeachae* infection, consistent with AM having a role as a replicative niche, as previously suggested using murine bone marrow-derived macrophages [3]. In line with this, AM supported efficient translocation of *L. longbeachae* T4SS effectors into the cytosol and promoted a higher bacterial load over other phagocytes. Dot/Icm T4SS effectors are known to be used by *L. pneumophila* and *L. longbeachae* to establish the *Legionella*-containing vacuole [5] and promote bacterial replication in mice [3].

In contrast to the role of AM evidenced here, our data identified a protective role for neutrophils and MC. While MC protected later during the course of infection confirming previous data [12], neutrophils restricted *L. longbeachae* in the early stages of infection. The delayed function of MC in reducing bacterial load was consistent with a delayed infiltration and bacterial uptake compared tto neutrophils. Furthermore, MC were the only cell types positive for *L. longbeachae* at later stages of infection. Neutrophils were previously suggested to be protective as they strongly infiltrate the lung during infection [3,42]. Here, we provide direct evidence of their protective role by quantifying a strong increase in *L. longbeachae* CFU following neutrophil depletion. Our data on *L. longbeachae* are consistent with the protective role of neutrophils during *L. pneumophila* infection [13]. Further investigation is required to elucidate how MC promoted defence against *L. longbeachae* despite supporting translocation of bacterial effectors, albeit to a lower extent than AM.

A classic bactericidal mechanism of neutrophils is enhanced production of ROS e.g. via IFN-γ, which is a key cytokine in defence against both *L. longbeachae* [3] and *L. pneumophila* [10]. IL-18 was initially discovered as an IFN-γ-inducing factor [36]. Here, we could show that this pathway is crucial for defence against *L. longbeachae* infection, and that it supports the function of neutrophils and MC to restrict *L. longbeachae*. This differentiates *L. longbeachae* infection from *L. pneumophila* infection, where IL-18 apparently has no role and IL-12 drives protective IFN-γ responses [10]. Although IL-12 is also important for defence against *L. longbeachae* [3], IL-18 plays as an additional role to generate protective IFN-γ levels. Our results suggest that during *L. longbeachae* infection, IFN-γ secretion is regulated via the IL-18R on either NK cells or T cells. Redundant protective functions of NK cells and T cells have been demonstrated for defence against other bacterial pathogens including *Mycobacterium tuberculosis* [46]. These results further highlight the differences between *L. longbeachae* and *L. pneumophila* as for the latter it has been shown that depletion of NK cells alone impaired bacterial clearance [47].

We have identified a large network of bronchiolar ciEpC expressing IL-18R1 that was, nevertheless, not required for protection against *L. longbeachae*. IL-18 apparently did not modulate ciliary beating frequency or mucus production, two key processes for mucociliary clearance and, accordingly, IL-18R1 expression by EpC in the lung did not confer protection against *L. longbeachae*. IL-18R expression by intestinal epithelial cells has been shown to promote inflammation, goblet cell death, and loss of barrier integrity during colitis [48]. Further investigations are required to understand its function in bronchiolar ciliated EpC.

In summary, our study provides an insight into the innate immune response to *L. longbeachae* infection, which is an understudied cause of life-threatening respiratory infection in humans. We show in a mouse model that infiltrating neutrophils and MC restrict bacterial infection while resident AM potentiate infection. Additionally, we define the importance of the IL-18–IFN-γ–ROS axis during infection, which may support the development of future therapeutic strategies against Legionellosis.

## Acknowledgements

We thank Christine Schmidt and Annika Völkel for excellent technical assistance. We would like to thank the Flow Cytometry Core Facility of the Medical Faculty at the University of Bonn for providing support and instrumentation funded by the Deutsche Forschungsgemeischaft (DFG, German Research Foundation) project-IDs: 387333827, 216372545, 216372401, 387335189, 471514137, 389568007. We thank Jennifer Dietrich and Prof. Daniela Wenzel for their excellent support in the establishment of ALI cultures. We acknowledge the HET and iFET facilities of the Medical Faculty, University of Bonn as well as the Biological Research Facility, Bio21 Institute for Biotechnology and Molecular Science for their animal husbandry. We thank the Melbourne Cytometry Platform (Bio21 node) for provision of cytometry services.

Work in the laboratory of N. Garbi is supported by DFG grants IRTG2168 – project-ID 272482170; SFBTRR237 – project-ID 369799452; SFB1454 – project-ID 432325352; Excellence Strategy EXC2151 – project-ID 390873048. Work in the laboratories of I. R. van Driel and E. L. Hartland was supported by the Victorian Government’s Operational Infrastructure Support Program and awards from the University of Melbourne and the Australian National Health and Medical Research Council (NHMRC), including APP1175976. M.F. was supported by a Graduate Research Scholarship from the University of Melbourne. Work in the laboratory of D. Wachten is supported by the DFG: SFB 1454 – Project-ID 432325352, TRR333/1 – Project-ID 450149205, FOR5547 - Project-ID 503306912, WA 3382/8-1 – Project-ID 513767027, under Germany’s Excellence Strategy – EXC2151 – Project-ID 390873048, and the Else Kröner Fresenius Foundation (2021.EKFSE.53). JNH was supported with a PhD fellowship from the Boehringer Ingelheim Fonds. We thank Dr. Luis Alvarez (caesar, Bonn) for supporting high-speed imaging equipment for imaging the ciliary beating in lung slices, in particular, the PCO Dimax high-speed camera.

## Author contributions

L.O., V.M.S., H.L.R., E.L.H., J.N.H., M.J.F., N.K., D.P., D.W., W.K., A.S.B., and G.N., participated in experimental design, planning, methodology, investigation, analysis, and/or visualization. E.L.H, I.R.v.D., G.N., and N.G., performed the conceptualization, funding acquisition, supervision, project administration. L.O., I.R.v.D., E.L.H., and N.G. drafted the manuscript. All authors contributed to the interpretation of data and approved the final version of the manuscript.

## Competing interests

The authors declare no competing interests.

## Abbreviations

AM: Alveolar macrophages
CBF: Ciliary beating frequency
CFU: Colony forming unit
ciEpC: Ciliated epithelial cells
HBEC: Human bronchiolar epithelial cells
ID: Infection dose
IFN: Interferon
IL: Interleukin
ILC: innate lymphoid cells
ImC: Immune cells
LOD: Limit of detection
L. lo: Legionella longbeachae
MC: Monocyte-derived cells
MFI: Mean fluorescence intensity
T4SS: Type IV secretion system
UMAP: Uniform Manifold Approximation and Projection

## Notes

### Competing Interest Statement

The authors have declared no competing interest.

