## Supplementary for "Opposing roles of resident and infiltrating immune cells in the defence against *Legionella longbeachae* via IL-18R/IFN-γ/ROS axis"

**Supplementary Table 1: Antibodies for flow cytometry**

| Antigen | Clone | Conjugate | Manufacturer |
|---|---|---|---|
| CD11b | M1/70 | BUV395 | BD Biosciences, Franklin Lakes, NJ, USA |
| | M1/70 | FITC | BD Biosciences, Franklin Lakes, NJ, USA |
| CD11c | N418 | PB | BioLegend, San Diego, CA, USA |
| | N418 | FITC | BioLegend, San Diego, CA, USA |
| CD19 | 6D5 | FITC | BioLegend, San Diego, CA, USA |
| | 6D5 | BV786 | BioLegend, San Diego, CA, USA |
| CD218a/ IL-18R1 | P3TUNYA | PE | Invitrogen, Thermo Fisher Scientific, Waltham, MA, USA |
| CD31 | MEC 13.3 | FITC | BioLegend, San Diego, CA, USA |
| CD326/ EpCAM | G8.8 | AF647 | BioLegend, San Diego, CA, USA |
| CD3ε | 145-2C11 | FITC | BioLegend, San Diego, CA, USA |
| | 17A2 | BV650 | BioLegend, San Diego, CA, USA |
| CD4 | GK1.5 | BUV737 | BD Biosciences, Franklin Lakes, NJ, USA |
| CD45 | 30-F11 | AF700 | BioLegend, San Diego, CA, USA |
| | 30-F11 | PO | Invitrogen, Thermo Fisher Scientific, Waltham, MA, USA |
| CD45.2 | 104 | AF488 | BioLegend, San Diego, CA, USA |
| | 104 | PB | BioLegend, San Diego, CA, USA |
| CD64 | X54-5/7.1 | PE | BioLegend, San Diego, CA, USA |
| | X54-5/7.1 | BV605 | BioLegend, San Diego, CA, USA |
| CD8a | 53-6.7 | BUV805 | BD Biosciences, Franklin Lakes, NJ, USA |
| Ly-6G | 1A8 | BV711 | BioLegend, San Diego, CA, USA |
| | 1A8 | PE | BioLegend, San Diego, CA, USA |
| | 1A8 | FITC | BD Biosciences, Franklin Lakes, NJ, USA |
| MHC-II | M5/114.15.2 | PE/Cy7 | BioLegend, San Diego, CA, USA |
| NK1.1 | PK136 | FITC | BioLegend, San Diego, CA, USA |
| | PK136 | AF647 | BioLegend, San Diego, CA, USA |
| SiglecF | E-50-2440 | AF647 | BD Biosciences, Franklin Lakes, NJ, USA |
| | E-50-2440 | PE | BD Biosciences, Franklin Lakes, NJ, USA |
| TCRβ | H57-597 | AF700 | BioLegend, San Diego, CA, USA |
| TCRγ/δ | GL3 | PE/Cy7 | BioLegend, San Diego, CA, USA |
| Ter119 | TER-119 | FITC | BioLegend, San Diego, CA, USA |
| Thy1.2 | 30-H12 | BV785 | BioLegend, San Diego, CA, USA |
| | 53-2.1 | PB | BioLegend, San Diego, CA, USA |

**Supplementary Table 2: Antibodies for confocal microscopy**

| Antigen | Clone | Conjugate | Manufacturer |
|---|---|---|---|
| Aquaporin5 | Rabbit polyclonal | Purified | Abcam, Cambridge, UK |
| CD45 | 30-F11 | AF594 | BioLegend, San Diego, CA, USA |
| EpCAM | G8.8 | AF647 | BioLegend, San Diego, CA, USA |
| EpCAM | G8.8 | AF488 | BioLegend, San Diego, CA, USA |
| Uteroglobin | Rabbit polyclonal | Purified | Abcam, Cambridge, UK |
| Acetylated tubuilin IV | Mouse IgG2b 6-11B-1 | Purified | Sigma-Aldrich, St. Louis, MO, USA |
| Rabbit | Donkey polyclonal | DL405 | Jackson ImmunoResearch Laboratories, West Grove, PA, USA |
| Rabbit | Donkey polyclonal | AF647 | Jackson ImmunoResearch Laboratories, West Grove, PA, USA |
| Mouse | Donkey polyclonal | DL405 | Jackson ImmunoResearch Laboratories, West Grove, PA, USA |

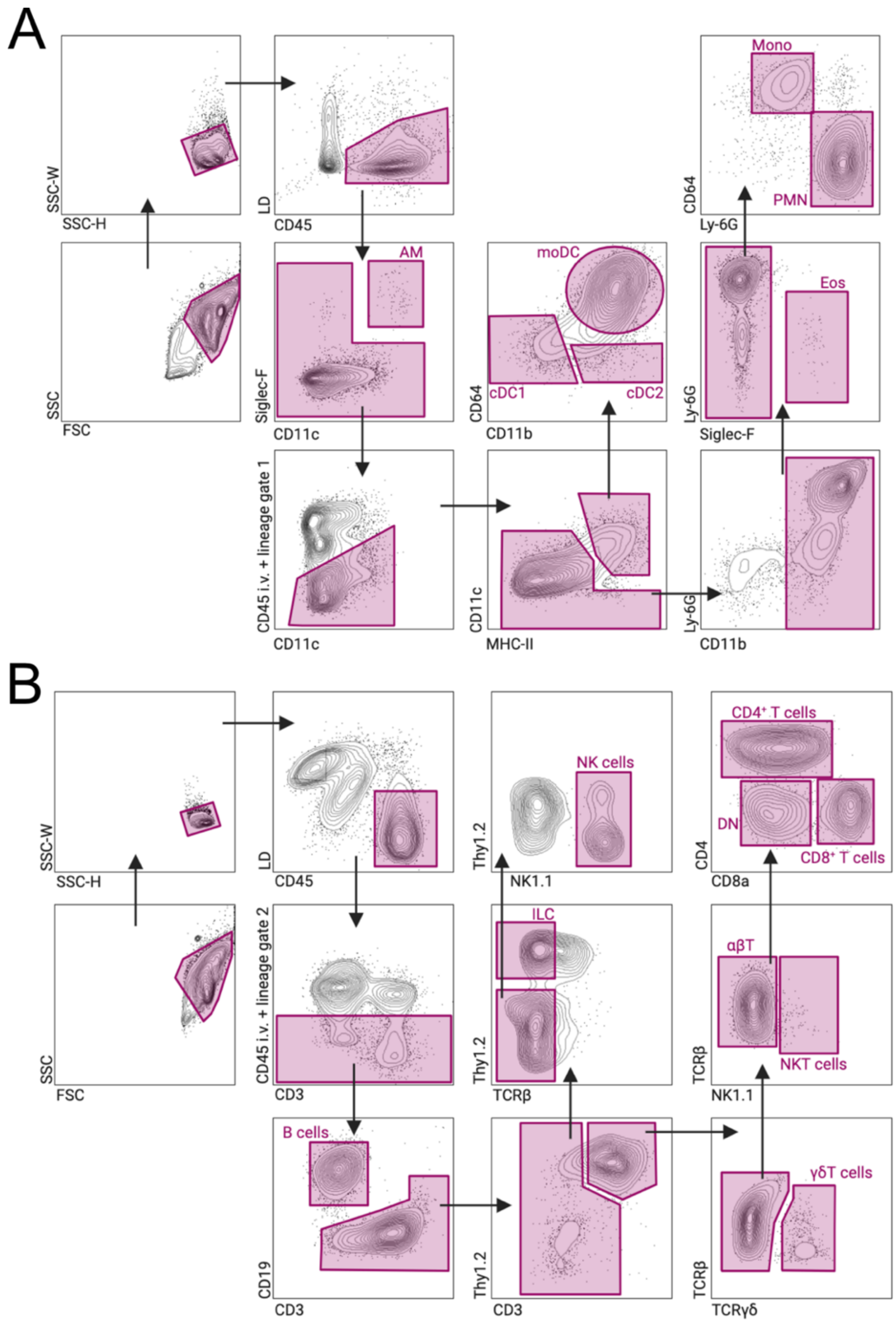


**Supplementary Figure 1: Gating strategy to identify myeloid (A) and lymphoid (B) cell populations using multi-color flow cytometry.** Pulmonary single cell-suspensions were analysed by multi-color flow cytometry. **(A)** Lineage 1: CD3, CD19, NK1.1, and Ter119. Alveolar macrophages (AM), Live CD45$^{+}$ Siglec-F$^{+}$ CD11c$^{+}$. All other cell populations were gated on Live CD45$^{+}$CD45i.v.$^{-}$Lin1$^{-}$: Dendritic cells (DC), MHC-II$^{+}$CD11c$^{+}$. Monocyte-derived Dendritic cells (moDC), CD11b$^{+}$CD64$^{+}$. Conventional Dendritic cells 1 (cDC1), CD11b$^{-}$CD64$^{-}$ DC. Conventional Dendritic cells 2 (cDC2), CD11b$^{+}$CD64$^{-}$ DC. Eosinophils (Eos), CD11c$^{-}$ CD11b$^{+}$ Siglec-F$^{+}$. Neutrophils (PMN), CD11c$^{-}$CD11b$^{+}$Siglec-F$^{-}$Ly-6G$^{+}$CD64$^{-}$. Monocytes (Mono), CD11c$^{-}$CD11b$^{+}$Siglec-F$^{-}$Ly-6G$^{-}$ CD64$^{+}$. **(B)** Lineage 2: CD11c, Ly-6G, and Ter119. Lymphoid cells (Live CD45$^{+}$ CD45i.v.$^{-}$ Lin2$^{-}$) were subdivided into: B cells, CD3$^{-}$ CD19$^{+}$. Innate lymphoid cells (ILC), CD19$^{-}$ CD3$^{-}$ Thy1.2$^{+}$. NK cells, CD19$^{-}$ CD3$^{-}$ Thy1.2$^{-}$ NK1.1$^{+}$. NKT cells, CD19$^{-}$ CD3$^{+}$ Thy1.2$^{+}$ TCR$\gamma\delta^{-}$ NK1.1$^{+}$. $\alpha\beta$T cells, CD19$^{-}$ CD3$^{+}$ Thy1.2$^{+}$ TCR$\gamma\delta^{-}$ NK1.1$^{-}$ TCR$\beta^{+}$. CD4$^{+}$ $\alpha\beta$T cells, CD4$^{+}$ CD8$^{-}$. CD8$^{+}$ $\alpha\beta$T cells, CD4$^{-}$ CD8$^{+}$. $\gamma\delta$T cells, CD19$^{-}$ CD3$^{+}$ Thy1.2$^{+}$ TCR$\gamma\delta^{+}$.

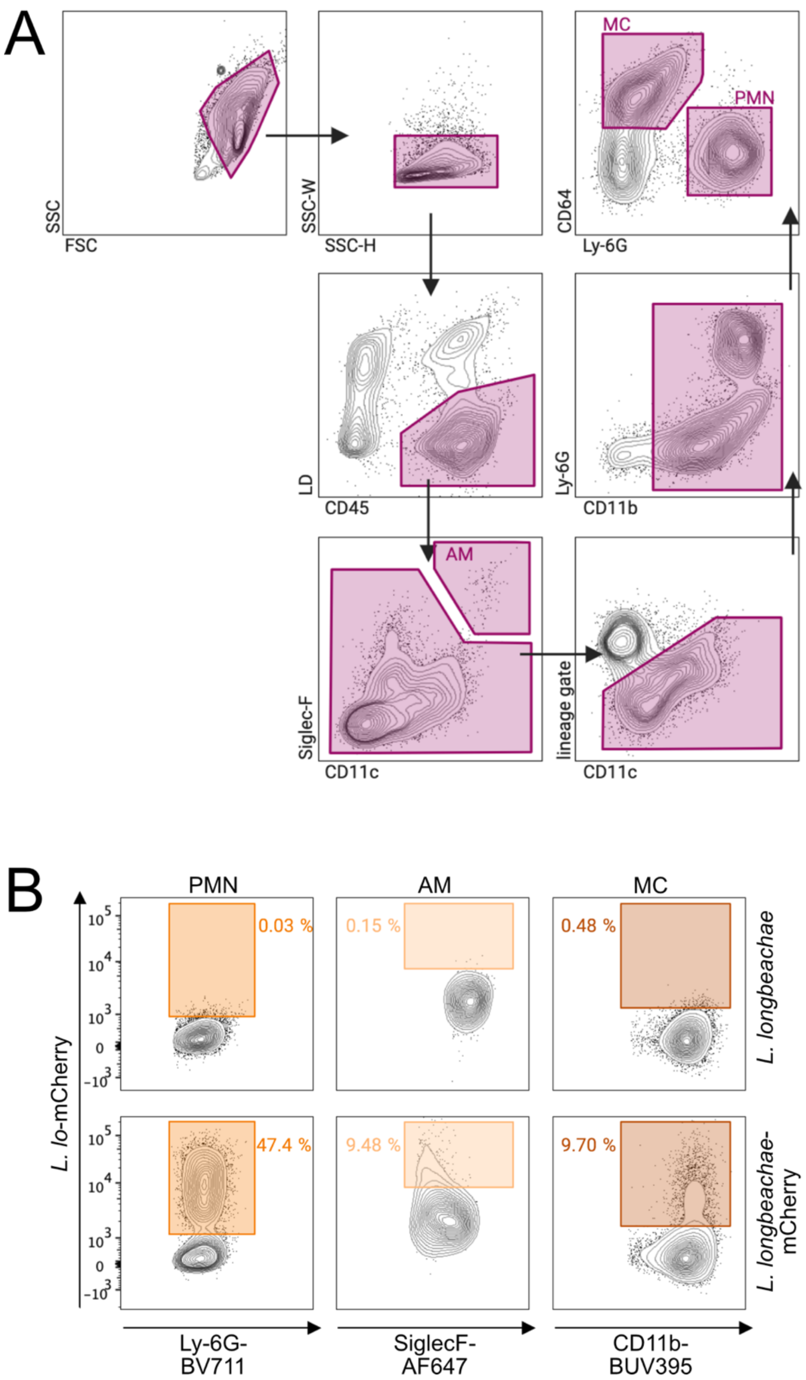


**Supplementary Figure 2: *L. longbeachae* locates in neutrophils (PMN), Alveolar macrophages (AM) and monocytes/monocyte-derived cells (MC).** C57BL/6J mice were infected i.n. with $10^4$ L. lo-mCherry CFU. Pulmonary single cell-suspensions were analysed by multi-color flow cytometry. **(A)** Single cells were gated by standard gating using SSC-H and SSC-W. Lineage exclusion gate included: CD3, CD19, NK1.1, and Ter119. Alveolar macrophages (AM), Live $CD45^+$ Siglec-$F^+$ $CD11c^+$. Neutrophils (PMN), Live $CD45^+$ $Lin^-CD11b^+$ Ly-$6G^+$ $CD64^-$. Monocyte-derived cells (MC), Live $CD45^+$ $Lin^-$ $CD11b^+$ Ly-$6G^-$ $CD64^+$. **(B)** Representative flow cytometry plots of indicated phagocyte populations 3 days after infection with WT L. lo (top panel) or L. lo-mCherry (bottom panel).

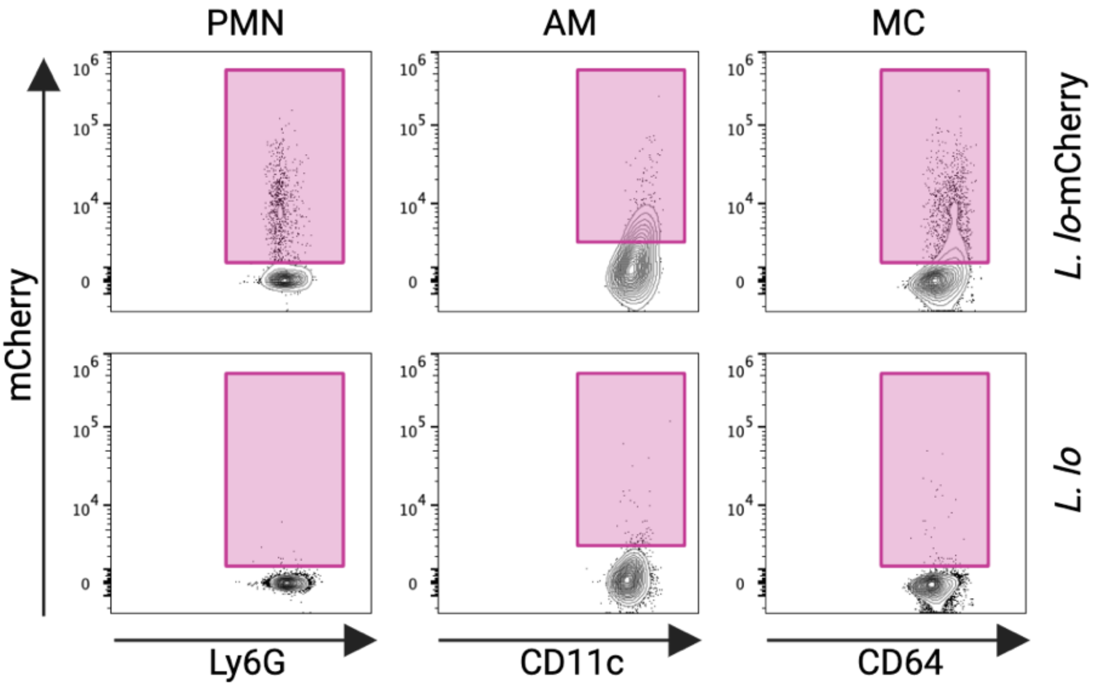


**Supplementary Figure 3: Spectral flow cytometry of the indicated phagocytes from infected mice.** ***L. longbeachae*-mCherry-containing cells from the lung of mice infected as indicated** identified using a 5-laser SONY ID7000 spectral flow cytometer. Autofluorescence was removed by spectral unmixing. Gating of cells as shown in **Figure S2**.

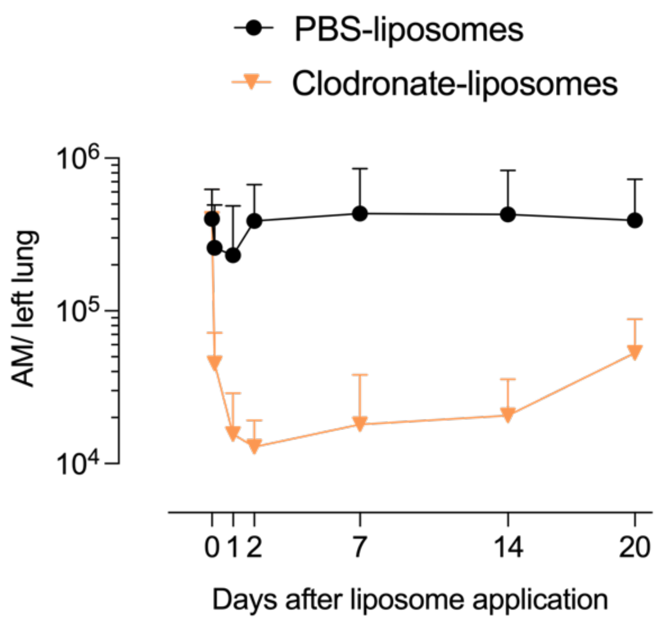


**Supplementary Figure S4: Long-term depletion of AM using clodronate-loaded liposomes.** C57BL/6J mice received 50 µl clodronate- or PBS-loaded liposomes i.t. and AM were quantified over time from lung cell suspension by flow cytometry.

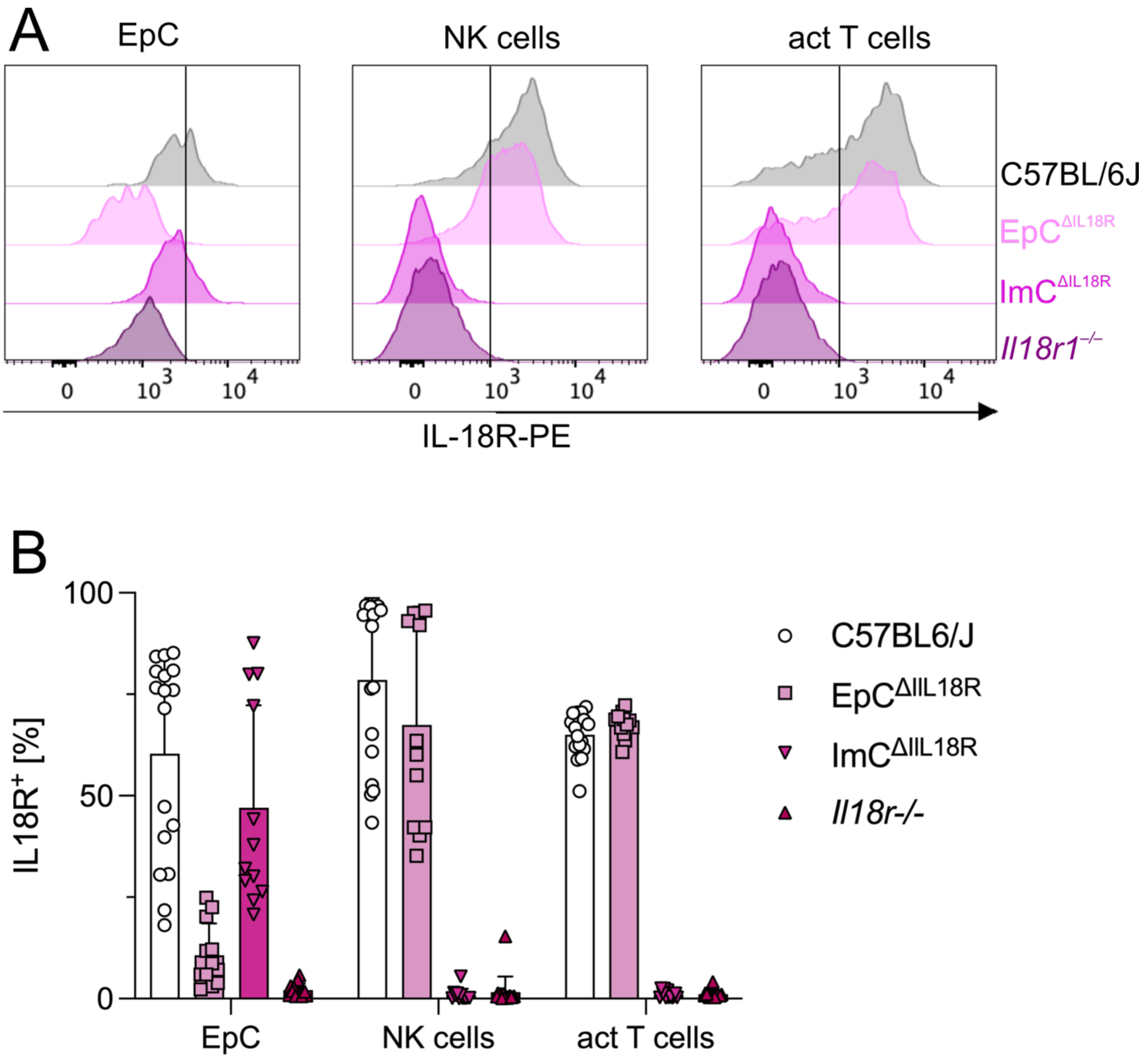


**Supplementary Figure 5: IL-18R1-depletion efficiency in EpC$^{\Delta IL18R}$ and ImC$^{\Delta IL18R}$ mice.** C57BL/6J, EpC$^{\Delta IL18R}$, ImC$^{\Delta IL18R}$, and constitutive *Il18r1*$^{-/-}$ mice were intranasally (i.n.) infected with 5x $10^4$ *L. longbeachae* (*L. lo)* CFU. IL-18R1 expression on the indicated lung cells was quantified 5 days after infection. **(A)** Representative flow cytometry histograms. **(B)** quantification of IL-18R1$^+$ cells from (A). Each symbol represents an individual mouse. EpC, epithelial cells; act, activated T cells (CD44$^{hi}$). Data are pooled from 2

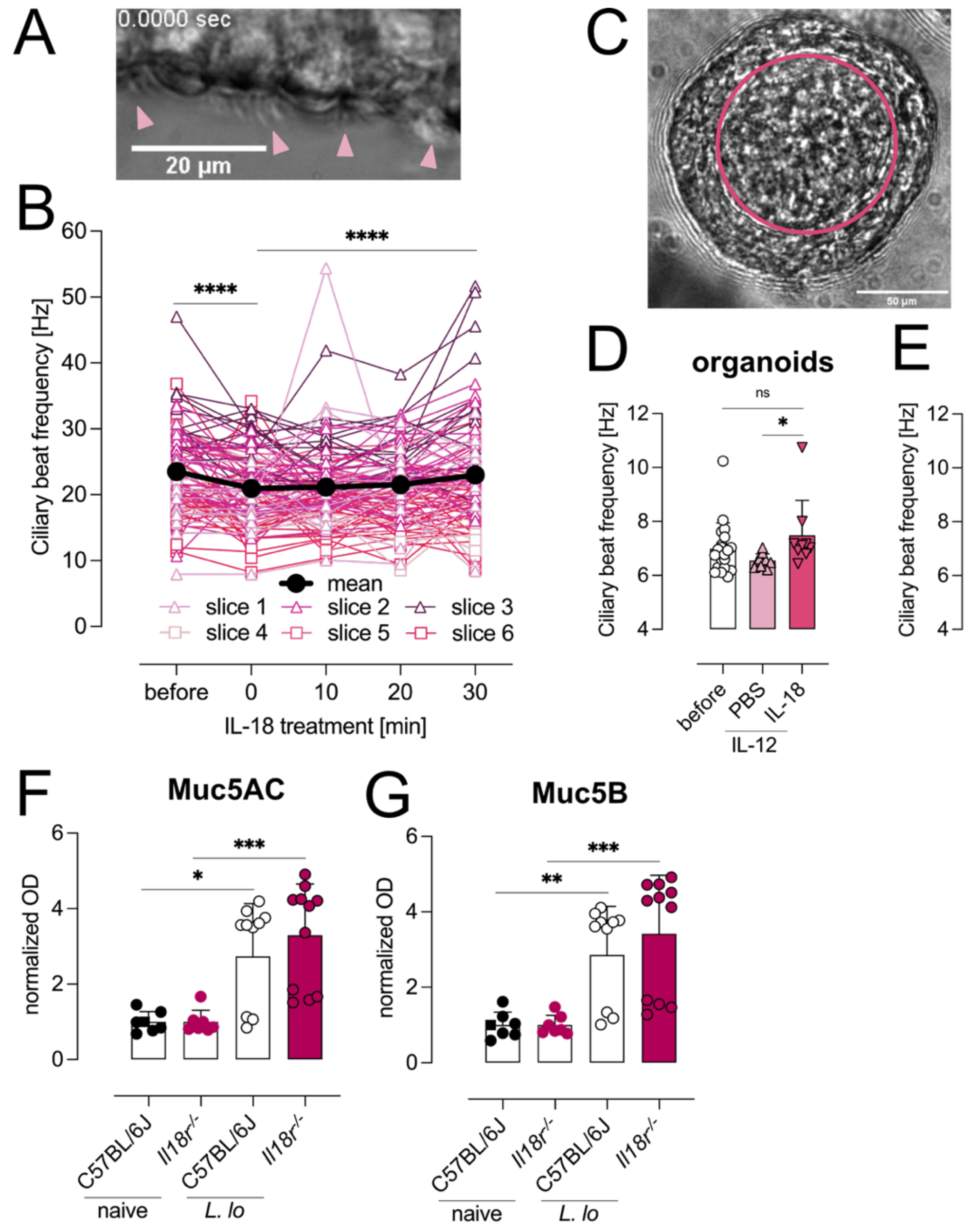


**Supplementary Figure 6: IL-18 does not influence ciliary beating frequency or mucus production. (A, B)** Quantification of ciliary beating frequency (CBF) in bronchiolar EpC using *ex vivo* lung vibratome sections. (A) Frame showing ciliated EpC (arrow heads) obtained from **Supplementary Video 2**. (B) CBF quantification from lung vibratome slices as shown in (A) using highspeed video microscopy before and during 30 min treatment with rIL-18. Symbol shape represents different mice (n=2, 3 slices per mouse). **(C, D)** CBF quantification in bronchiolar EpC using primary human bronchiolar epithelial cells grown as organoids. (C) Frame showing ciliary beating (within circle) obtained from **Supplementary Video 3**. (D) CBF in organoids quantified 16h after treatment as indicated. **(E)** CBF quantification in bronchiolar EpC using primary human bronchiolar epithelial cells grown as ALI cultures. Cultures were treated as in (D). **(F, G)** Quantification of mucus proteins MUC5AC (F) and MUC5B (G) in the lung homogenate 3 days after i.n. infection of the indicated mouse strains with 5x $10^4$ *L. lo*. * $P < 0.05$; ** $P < 0.01$; *** $< 0.001$, **** $P < 0.0001$ (Mann-Whitney test (F, G) or repeated measures one-way ANOVA (B, E) or Kruskal-Wallis (D) test with Turkey's or Dunn's multiple comparisons tests). (B) data are pooled from two experiments with two mice (3 lung slices per mouse). (D, E) data are pooled from two (D) experiments or from one experiment (E) with symbols representing single wells. (F, G) data are pooled from two experiments with symbols representing single mice.

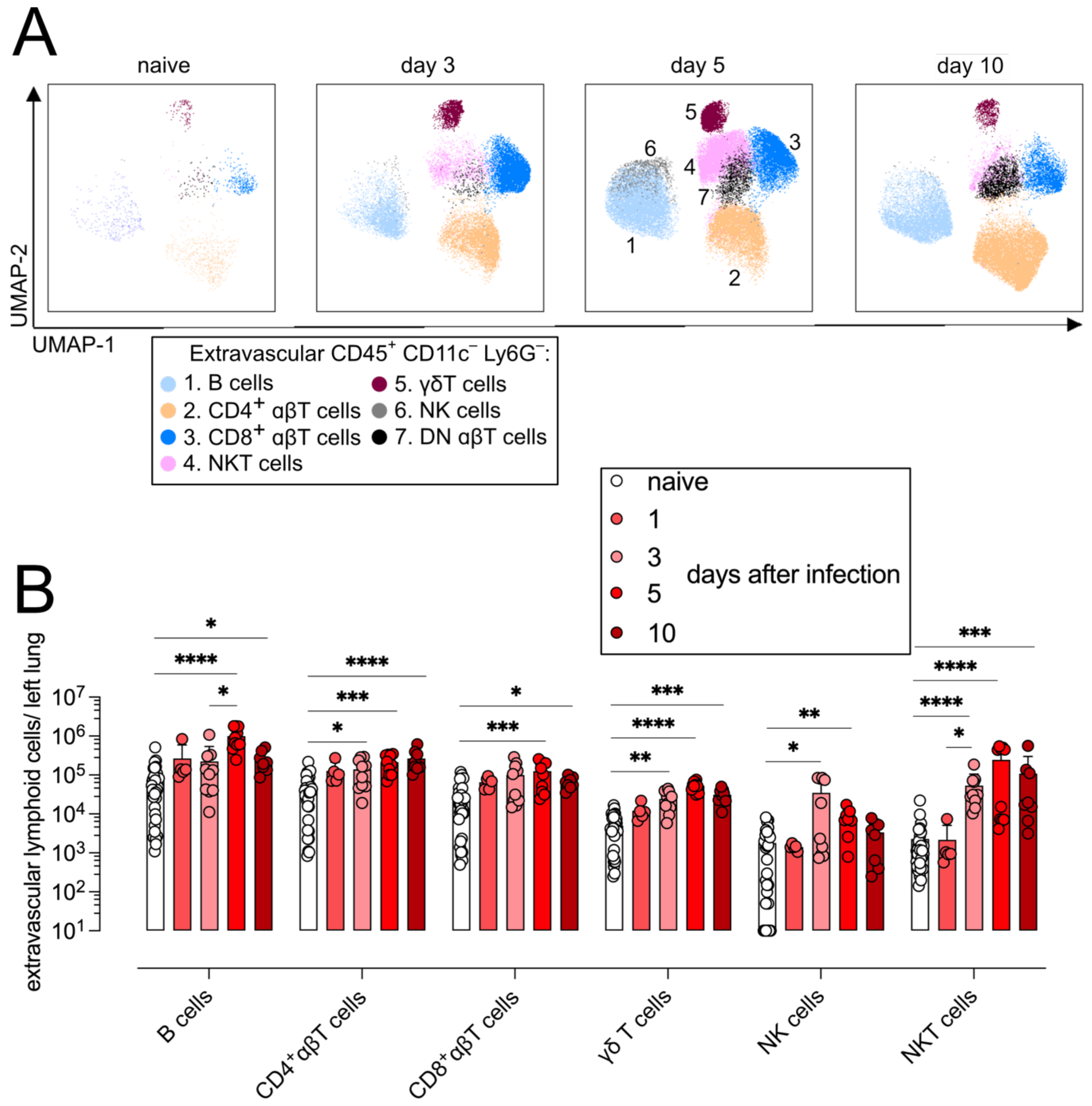


**Supplementary Figure 7: Quantification of lymphoid cells during infection.** C57BL/6J mice were intranasally (i.n.) infected with $10^4$ *L. longbeachae* (*L. lo)* CFU. **(A)** Flow cytometry UMAP plots of extravascular CD19⁻ NK1.1⁻ CD3⁻ myeloid cells at the indicated time points. Gating strategy to identify specific immune cell populations is shown in Fig S1. Lymphoid populations are indicated in day 5 plot. **(B)** Quantification of indicated immune cells from (A). Symbols represent single mice. moDC = monocyte-derived dendritic cells, cDC = conventional dendritic cells. Kruskal-Wallis test (with Dunn's multiple comparisons test for inter-group differences) was performed for statistical analysis. Data are shown as mean +SD. * $P < 0.05$; ** $P < 0.01$; *** $P < 0.001$; **** $P < 0.0001$.

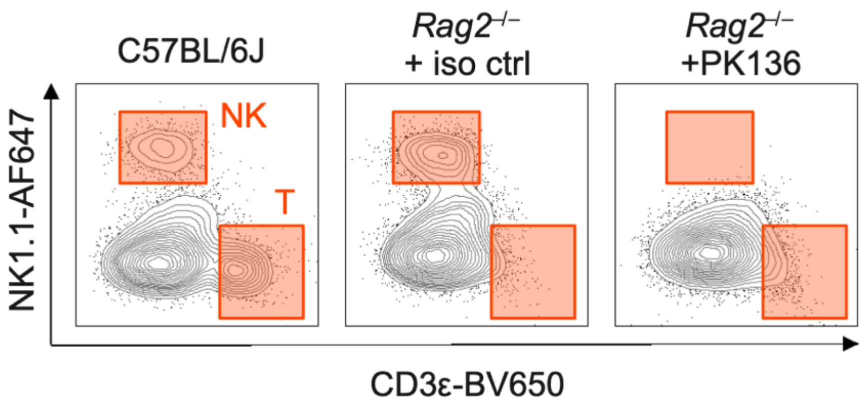


**Supplementary Figure 8: NK cell depletion in *Rag2*$^{-/-}$ mice.** *Rag2*$^{-/-}$ mice treated with depleting anti-NK.1. antibody (PK136) or isotype control (iso ctrl, rIgG2a) and i.n. infected with 5x $10^4$ *L. lo*-mCherry CFU. C57BL/6J mice were i.n. infected with 5x $10^4$ *L. lo* CFU. Representative flow cytometry plot showing lack of NK cells and T cells in PK136 treated *Rag2*$^{-/-}$ mice compared to the WT and isotype control on day 5 after infection.

**Supplementary Material and Methods:**

Organotypic cell culture of human bronchiolar epithelial cells:

Human bronchiolar epithelial cells (HBEC) were cultivated at 37 °C and 5 % $CO_2$ under humidified conditions in Basal medium (PneumaCult$^{TM}$-ExPlus Basal medium + 1x PneumaCult$^{TM}$-ExPlus 50X Supplement + 0.2x Hydrocortisone + 100 U/ ml Penicillin-Streptomycin). To differentiated HBEC in an ALI culture or organoid system the PneumaCult$^{TM}$ ALI kit (StemCell, Canada) or the PneumaCult$^{TM}$ Airway Organoid kit (StemCell, Canada) were used, respectively, according to the manufacturer's instructions.

Briefly, to generate ALI cultures 8 x $10^4$ HBEC per well were seeded in collagen-coated trans-well inserts in a 12 well-plate. Cells were grown in Basal medium for 5 days before an air stimulus was induced and medium changed to ALI Maintenance medium (StemCell, Canada; PneumaCult$^{TM}$-ALI Basal medium + 1x PneumaCult$^{TM}$-ALI 10X supplement + PneumaCult$^{TM}$-ALI 100X Maintenance Supplement 50x + 0.00032 % Heparin + 0.8x Hydrocortisone + 100 U/ ml Penicillin-Streptomycin) for 21 days.

To generate bronchiolar organoids, 2x $10^3$ HBEC were seeded in a 50 µl Matrigel (Corning Inc., USA) dome per well in 24 well plates. 5 days after seeding the Organoid Seeding medium (PneumaCult$^{TM}$-Organoid Basal medium + 1x PneumaCult$^{TM}$-Organoid Seeding Supplement + 100 U/ ml Penicillin-Streptomycin) was exchanged to Organoid Differentiation medium (PneumaCult$^{TM}$-Organoid Basal medium + 1x PneumaCult$^{TM}$-Organoid Differentiation Supplement + 0.00032 % Heparin + 0.8x Hydrocortisone + 100 U/ ml Penicillin-Streptomycin) for further 28 days.

Quantification of lung ciliary beating frequency:

Ciliary beating frequency (CBF) was quantified in *ex vivo* lung vibratome slices and *in vitro* in organotypically grown HBEC. To quantify CBF *ex vivo*, 10 µg LPS (Serotype 026:B6; Sigma-Aldrich, US) was i.t. administered to mice one day prior to sampling to upregulate IL-18Rβ expression. Lung vibratome slices were prepared as described under *Confocal microscopy*. Microphotographs of lung vibratome sections were recorded at 3 000 frames / sec using a high-speed video camera and subsequently treated with 100 ng/ ml rIL-18 for a period of 30 min. CBF was recorded every 10 min.

CBF was recorded *in vitro* from ALI cultures and organoids with 800 frames/ sec before and after overnight treatment with 100 ng/ ml IL-12 ± 100 ng/ ml IL-18. IL-12 was added to upregulate IL-18Rβ.

Image recordings were performed at 37°C using an inverted IX71 Olympus microscope equipped with a halogen lamp for illumination, a condenser, a phase ring blend, a 40x objective (NA 0.95, UPlanSApo, Olympus), a microscope incubator (Life Imaging Services, Basel, Switzerland), and a high-speed camera (PCO Dimax, PCO, Germany, for Vibratome slices; ORCA-Flash4.0 V3, C13220–20 CU, Hamamatsu, Japan, for the ALI cultures and organoids).

CBF was quantified in ImageJ/ Fiji (v2.8.0/1.53t) as described in [1]. All recorded images were concatenated to one video and the video was registered via the MultiStackReg plugin by Brad Busse (http://bradbusse.net/downloads.html) to eliminate artefacts resulting from whole-sample motion. The MultiStackReg plugin is an adaptation of another ImageJ plugin called "TurboReg" (http://bigwww.epfl.ch/thevenaz/turboreg/) [2]. After registration, the image was manually split info multiple smaller images with each image displaying one cell and its cilia. Each cell image was analysed with the custom-written ImageJ plugin Cilility_JNH (Version v0.0.1 for Vibratome slices; v0.2.1 for ALI cultures and organoids), which, after finishing the analysis of the data shown in this study, was refurbished with the aim to make the plugin applicable to diverse data sets, renamed to “FreQ”, and described in [1]. Source code and plugins for FreQ (and Cilility_JNH) are available in the FreQ github repository: https://github.com/hansenjn/FreQ. The source code of Cilility_JNH v0.0.1 corresponds to the source code published in the first FreQ github commit c426545 (accessible through https://github.com/hansenjn/FreQ/tree/c426545c0b051399686abb1c0f1e237a3205ba5c) except for a few code changes that do not affect the presented analysis result – the code in commit c426545 can fully reproduce the analysis of vibratome slices presented in this study. The source code and software for Cilility version v0.2.1 is available here: https://github.com/hansenjn/FreQ/releases/tag/v0.2.1. Cilility_JNH was applied under the following settings: sampling rate of 3000 Hz for Vibratome slices, 400 Hz for ALI cultures and organoids; sliding window size for smoothing power spectrum of 5 Hz (all images); 20.0 percent lower powers used for thresholding power spectrum (all images); 1.5-fold SD used for power thresholding (all images); max accepted frequency for filtering of 90.0 Hz (all images); min accepted frequency for filtering of 5.0 Hz (all images). In Cilility_JNH, for each image, a trained investigator selected a region of interest (ROI) circumscribing the beating cilia from an individual cell, and the CBF was quantified from the pixels only in that ROI. As CBF the Cilility_JNH output parameter “Average freq (Hz)” was used.

regions of interest (ROIs), each circumscribing the beating cilia from an individual cell, were selected by a trained observer and CBF was quantified for each region of interest using a custom written ImageJ plugin (Cilility_JNH, v0.0.1). The underlying method to determine the CBF has been described previously [3]. After finishing the experiments and analysis for the data shown in this study, the ImageJ plugin was refurbished with the aim to make the plugin applicable to diverse data sets and published under the name “FreQ” (https://github.com/hansenjn/FreQ).

Mucus quantification in the lung homogenate:

Muc5B and Muc5AC expression was quantified in lung homogenate using Colorimetric ELISA kits (Novus Biologicals) according to manufacturer's instructions. Briefly, left lungs were homogenized in 250 µl PBS using tubes with ceramic beads (Bertin Technologies) at 4,000 rpm for 1 min. The homogenate supernatant was obtained after centrifugation at 13,000 rpm and 4 °C for 5 min and was stored at -80 °C. If required samples were diluted in PBS. The provided 96 well-plated pre-coated with anti-Muc5B- or anti-Muc5AC-antibody was prepared with 100 µl of samples and incubated for 90 min at 37 °C. Supernatant was aspirated and 100 µl Biotinylated Detection antibody was incubated on the plate for 60 min at 37 °C. After decanting the supernatant, the plate was washed 4x with 350 µl Wash buffer. 100 µl HRP conjugate was incubated on the plate for 30 min at 37 °C. Supernatant was decanted and the plate was washed 5x with 350 µl Wash buffer. 90 µl of Substrate reagent was incubate on the plate for max. 15 min at 37 °C in the dark. The incubation time was adjusted to the actual colour change and the colour development was ceased by adding 50 µl Stopping Solution. The $OD_{450}$ was recorded at the Tecan Safire2.
